# A conserved regulatory program drives emergence of the lateral plate mesoderm

**DOI:** 10.1101/261115

**Authors:** Karin D. Prummel, Christopher Hess, Susan Nieuwenhuize, Hugo J. Parker, Katherine W. Rogers, Iryna Kozmikova, Claudia Racioppi, Eline C. Brombacher, Anna Czarkwiani, Dunja Knapp, Sibylle Burger, Elena Chiavacci, Gopi Shah, Alexa Burger, Jan Huisken, Maximina H. Yun, Lionel Christiaen, Zbynek Kozmik, Patrick Müller, Marianne Bronner, Robb Krumlauf, Christian Mosimann

**Author notes:** contributed equally to this work (Karin D. Prummel, Christopher Hess).

## Abstract

Cardiovascular lineages develop together with kidney, smooth muscle, and limb connective tissue progenitors from the lateral plate mesoderm (LPM). How the LPM initially emerges and how its downstream fates are molecularly interconnected remain unknown. Here, we isolated a pan-LPM enhancer in the zebrafish *draculin* (*drl*) gene that provides specific LPM reporter activity from early gastrulation. *In toto* live imaging and lineage tracing of *drl*-based reporters captured the dynamic LPM emergence as lineage-restricted mesendoderm field. The *drl* pan-LPM enhancer responds to the transcription factors EomesoderminA, FoxH1, and MixL1 that combined with Smad activity drive LPM emergence. We uncovered specific *drl* reporter activity in LPM-corresponding territories of several chordates including chicken, axolotl, lamprey, *Ciona*, and amphioxus, revealing a universal upstream LPM program. Altogether, our work provides a mechanistic framework for LPM emergence as defined progenitor field, possibly representing an ancient mesodermal cell state that predates the primordial vertebrate embryo.

## Introduction

Key cell fates and organ systems in vertebrates emerge from multipotent progenitors within the embryonic mesoderm. Following gastrulation, the vertebrate mesoderm has been classically described to partition into axial, paraxial, and ventro-lateral domains (Gilbert, 2000; Gurdon, 1995). The latter, referred to as lateral plate mesoderm (LPM), is composed of highly motile cells and is mainly defined by its position adjacent to the somite-forming paraxial mesoderm (Hatada and Stern, 1994). Transplantation and lineage tracing experiments in several species have established that the LPM contains progenitor cells of the circulatory system, smooth muscle lineages, the kidneys (in amniotes often demarcated as intermediate mesoderm), and the limb connective tissue anlagen (Chal and Pourquié, 2017; Takasato and Little, 2015). Several transcription factors including Hand1/2, Tbx5, Osr1, FoxF1, Prrx1, Mesp1, and Etv2 are expressed in LPM territories and play overlapping roles in cell fate determination (Chal and Pourquié, 2017; Davidson and Zon, 2004; Takasato and Little, 2015), albeit not always with an evolutionarily conserved function (Yabe et al., 2016). During segmentation, the LPM principally segregates into the anterior LPM (ALPM) and posterior LPM (PLPM), which further divides into dorsal and ventral domains (somatopleure and splanchnopleure, respectively) (Gilbert, 2000). Several studies have uncovered mechanisms that control the lateral-to-medial or anterior-posterior specification of LPM-descendant organ precursors (Chal and Pourquié, 2017; Helker et al., 2015; Tonegawa et al., 1997). In contrast, it remains incompletely understood how the LPM arises from an initial mesendodermal population that goes on to form distinct endodermal and mesodermal progenitors. This is in part due to the lack of tools and markers to track LPM emergence genetically during development. Further, whether the LPM initially emerges as morphogenetic field in a molecularly coherent unit or as a loosely connected assembly of progenitor cells remains unclear.

Assessing the evolutionary context by which the LPM emerged as a developmental entity also remains challenging, in particular in extant jawless vertebrates such as lamprey or chordate models that do not form the full spectrum of LPM derivatives. Ancestral gene-regulatory repertoires that control higher-order structures in vertebrates previously have been indicated for somatic muscle in lamprey (Kusakabe and Kuratani, 2007) or for the putative equivalents of cardiac and hematopoietic progenitors in amphioxus (Pascual-Anaya et al., 2013). Anterior-to-posterior expression domains of key LPM transcription factors including Tbx1/10 and Hand are conserved in lampreys and amphioxus (Onimaru et al., 2011). Furthermore, the tunicate *Ciona* forms cardiac lineages that display genetic regulatory circuits homologous to the cardiac LPM progenitors found in vertebrates (Kaplan et al., 2015; Racioppi et al., 2019). These observations suggest the existence of an ancient, defined regulatory program that delineates prospective LPM progenitors in a common chordate ancestor, dating back to the Cambrian explosion 520-540 million years ago.

Several mammalian *cis*-regulatory elements with broad LPM activity have been reported; these include a small upstream enhancer of mouse and human *HoxB6* (Becker et al., 1996), an upstream enhancer of mouse *Gata4* (Rojas et al., 2005), and a downstream enhancer of mouse *Bmp4* (Chandler et al., 2009). In line with a ventral LPM origin, the *Gata4* LPM enhancer responds to Smads downstream of BMP signaling (Rojas et al., 2005). Nonetheless, the activities driven by these enhancer elements in mice confine to the PLPM and not pan-LPM readouts. In zebrafish, the ventrally emerging LPM forms during somitogenesis into a patchwork of bilateral gene expression domains, including the conserved LPM genes *hand2*, *pax2.1*, *scl*, *lmo2*, *etv2*, and *tbx5* (Davidson and Zon, 2004). In contrast, transgenic reporters based on the 6.35 kb *cis*-regulatory region of the zebrafish-specific gene *draculin* (*drl*) selectively label the entire LPM from its emergence during gastrulation through initial differentiation (Mosimann et al., 2015). Cre/*lox*-mediated genetic lineage analysis has established that early *drl* reporter expression labels the LPM progenitors forming cardiovascular, blood, kidney, intestinal smooth muscles (iSMCs), and pectoral fin mesenchyme fates (Felker et al., 2016; Felker et al., 2018; Gays et al., 2017; Henninger et al., 2017; Mosimann et al., 2015). These observations suggest that the 6.35 kb *drl* region harbors *cis*-regulatory elements active throughout the prospective LPM starting from gastrulation, raising the possibility that these regulatory elements read out a hypothetical pan-LPM program.

Here, we dissected the 6.35 kb *drl cis*-regulatory elements and uncovered an intronic enhancer, +*2.0drl*, that is necessary and sufficient in zebrafish for driving LPM-specific expression in all presumptive LPM progenitors from early gastrulation to early somitogenesis. Panoramic SPIM and Cre/*lox*-mediated genetic lineage tracing of *drl* reporters demonstrated that the zebrafish LPM forms from a restricted mesendoderm territory during gastrulation. To uncover the upstream regulatory program read out by the +*2.0drl* pan-LPM enhancer, we combined ChIP data, Cas9-based crispant analysis, and reporter assays. This enabled us to identify the combination of mesendoderm transcription factors EomesA, FoxH1, and MixL1 as sufficient to drive pan-LPM activity. Our data suggest that the combined activity of these transcription factors demarcates a dedicated mesendoderm progenitor field during zebrafish gastrulation, restricting the emergence of LPM progenitors to their future lateral position. In cross-species assays, we observed specific activity of the zebrafish +*2.0drl* pan-LPM enhancer in LPM-corresponding territories in chicken, axolotl, lamprey, *Ciona*, and amphioxus embryos. These results demonstrate that the +*2.0drl* enhancer reads out a universal LPM progenitor program that is conserved across chordates, defining a core transcription factor code for LPM formation. Our data provide a developmental framework for charting the earliest emergence of LPM progenitors across chordates.

## Results

### The LPM emerges as a dedicated mesendoderm population

To resolve the dynamics of LPM emergence *in toto*, we performed time course experiments using single-plane illumination microscopy (SPIM) (Fig 1A-D) and panoramic projections (Schmid et al., 2013) (Fig 1E-H) of reporter-transgenic zebrafish embryos based on the full-length *drl cis*-regulatory region. *drl*:*EGFP*-expressing LPM precursors became detectable by early gastrula stages (50% epiboly) and continuously condensed along the embryo margin through the end of gastrulation (tailbud stage) (Fig 1A,B,F; Movie S1, S2). From tailbud stage onward, *drl:EGFP*-marked LPM formed a continuous band of cells with condensing anterior and posterior segments (Fig 1C,D,G,H). We confirmed that this EGFP-positive cell band encompasses the bilateral stripes of several established LPM sub-domain markers by comparing a series of overlapping expression domains from distinct reporter lines (Fig 1I-L). First, *lmo2:dsRED2* labels embryonic hematopoietic and vascular tissues (Zhu et al., 2005), and its expression overlaps with medial *drl:EGFP*-expressing cells in the ALPM and PLPM (Fig 1I). *scl:EGFP* (Zhang and Rodaway, 2007) also co-expressed with *drl:mCherry* in the most medial PLPM domain and in a small ALPM population (Fig 1J). We find that while *pax2.1:EGFP* in early somitogenesis marks midbrain-hindbrain precursor cells of ectodermal origin, the likewise *pax2.1:EGFP*-expressing PLPM-derived pronephric epithelial precursors (Picker et al., 2002) also express *drl:mCherry* (Fig 1K). Moreover, *hand2:EGFP* expression, which demarcates the lateral-most PLPM domain plus parts of the ALPM-derived heart field and pectoral fin precursors (Perens et al., 2016; Yelon et al., 2000), was also fully situated within the pan-LPM expression domain of *drl:mCherry* (Fig 1L). Taken together, these data provide a continuous view of the emerging LPM stripes from gastrulation in zebrafish and document that the LPM emerges around the entire circumference of the zebrafish embryo (Fig 1M).

**Figure 1:**
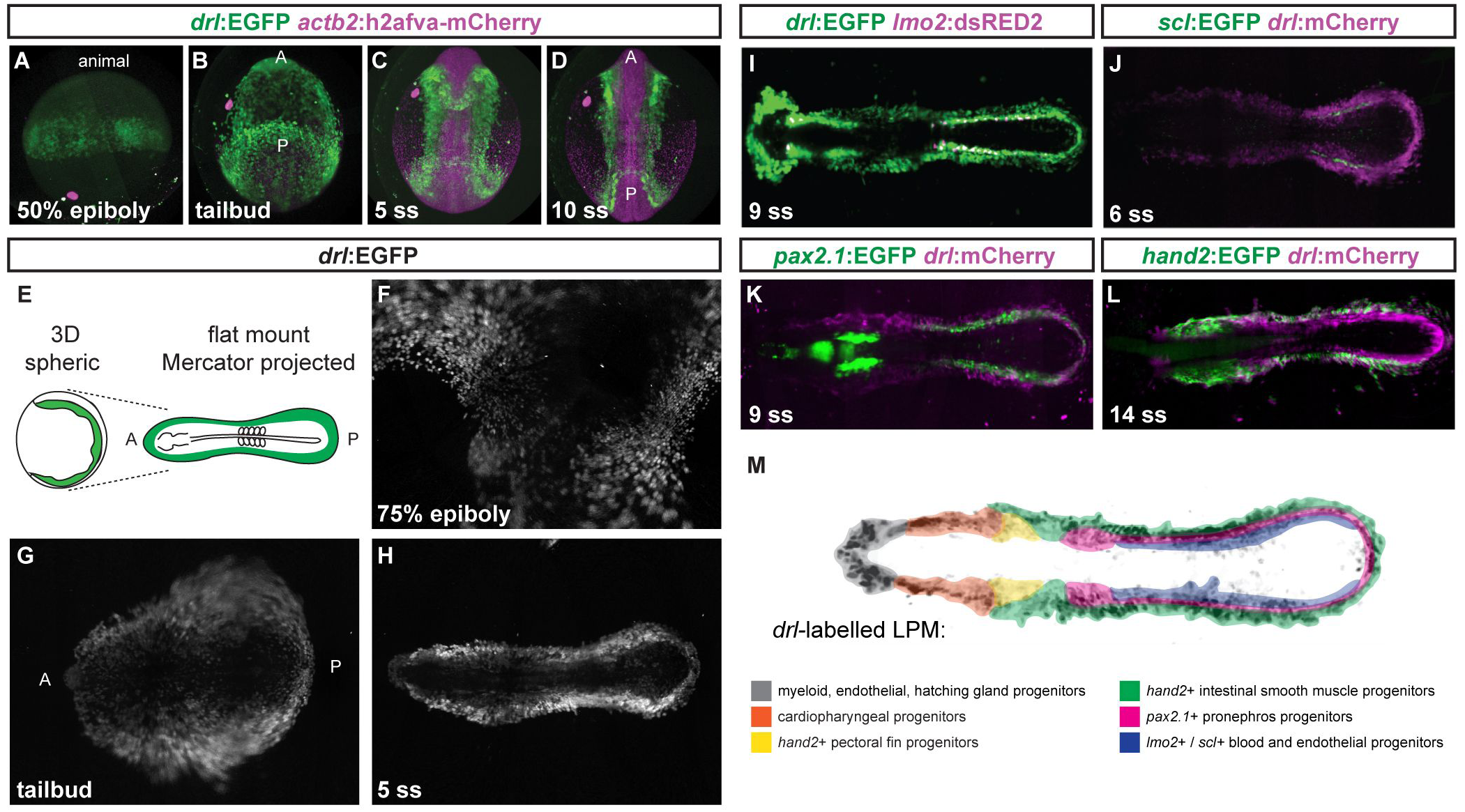
The LPM forms as a continuous field around the entire circumference of the developing zebrafish embryo. (**A-D**) Panoramic SPIM imaging of 50% epiboly to 10 ss embryos transgenic for *drl*:*EGFP* (green) and *actb2:h2afva-mCherry* (magenta); maximum-intensity-projected, lateral view (**A**), dorso-ventral views (anterior (a) to the top, posterior (p) bottom) (**B-D**). (**E-H**) Panoramic SPIM imaged *drl:EGFP* zebrafish embryos shown as 2D Mercator projections. (**E**) Schematic of Mercator projection of spherical embryo, anterior to the left; (**F-H**) Mercator projections at 75% epiboly, tailbud, and 5 ss stages. (**I-L**) Single time point projections (anterior to the left) of double-transgenic embryos for *drl* reporters and (**J**) *lmo2*, (**K**) *scl*, (**L**) *pax2.1*, or (**M**) *hand2* reporters co-expressed in dedicated LPM territories. (**M**) Summary schematic of the LPM fate territories partitioning during early somitogenesis in zebrafish.

We next sought to capture how the *drl*-expressing LPM emerges relative to the endoderm. Panoramic SPIM of the *sox17:EGFP*-positive endoderm reporter together with *drl:mCherry* revealed a population of double-positive cells from the onset of reporter detection through late gastrulation (Fig 2A-D; Fig S1A-D). After gastrulation, we detected a continuous band of *drl* reporter-positive cells around the developing embryo that was separated from the more medial endodermal *sox17* expression domain (Fig 2D; Movie S3, S4). To confirm whether endoderm progenitors are also marked by the *drl* reporter during gastrulation, we performed *drl:creERT2*-based genetic lineage tracing with the ubiquitous *hsp70l:loxP-STOP-loxP-EGFP* (*hsp70l:Switch*) and 4-OHT-based CreERT2 induction at discrete time points ranging from shield to 5-6 somite stages (ss) followed by analysis of labeling patterns at 72 hpf (Fig 2E-J). 4-OHT induction at shield stage marked LPM lineages including blood, endothelium, kidney, and iSMCs as the only mesodermal fates (Fig 2F-H) (Gays et al., 2017; Mosimann et al., 2015), while lineage labeling also marked broad territories within endodermal organs, including pancreas, liver, and pharynx/gut epithelium (Fig 2H; Fig S1A). 4-OHT induction at increasingly later time points gradually decreased endoderm labeling, with minimal to no endodermal lineage signals following 4-OHT induction at 5-6 ss (Fig 2I,J, Fig S1B, Fig S2B,C). In contrast, LPM structures remained robustly labeled as the exclusive mesoderm fate, consistent with previous work (Felker et al., 2018; Gays et al., 2017; Henninger et al., 2017; Mosimann et al., 2015). Additionally, we observed that the spatio-temporal contribution of *drl* reporter-expressing progenitors to endoderm differs along the anterior-posterior axis. We divided the embryo into four non-overlapping regions along the anterior-to-posterior axis (region I – IV) (Fig. S2A) and quantified the switching efficiency. The amount of lineage-labeled gut endothelium increased within individual embryos from the anteriorly located pharynx (region I) towards the caudal gut (region IV), independent of the stage of 4-OHT administration (Fig S2B,C). These results indicate that progenitors expressing the *drl* reporter become progressively restricted to an LPM fate from anterior to posterior with ongoing development, until by early somitogenesis *drl* reporter expression labels only early LPM.

**Figure 2:**
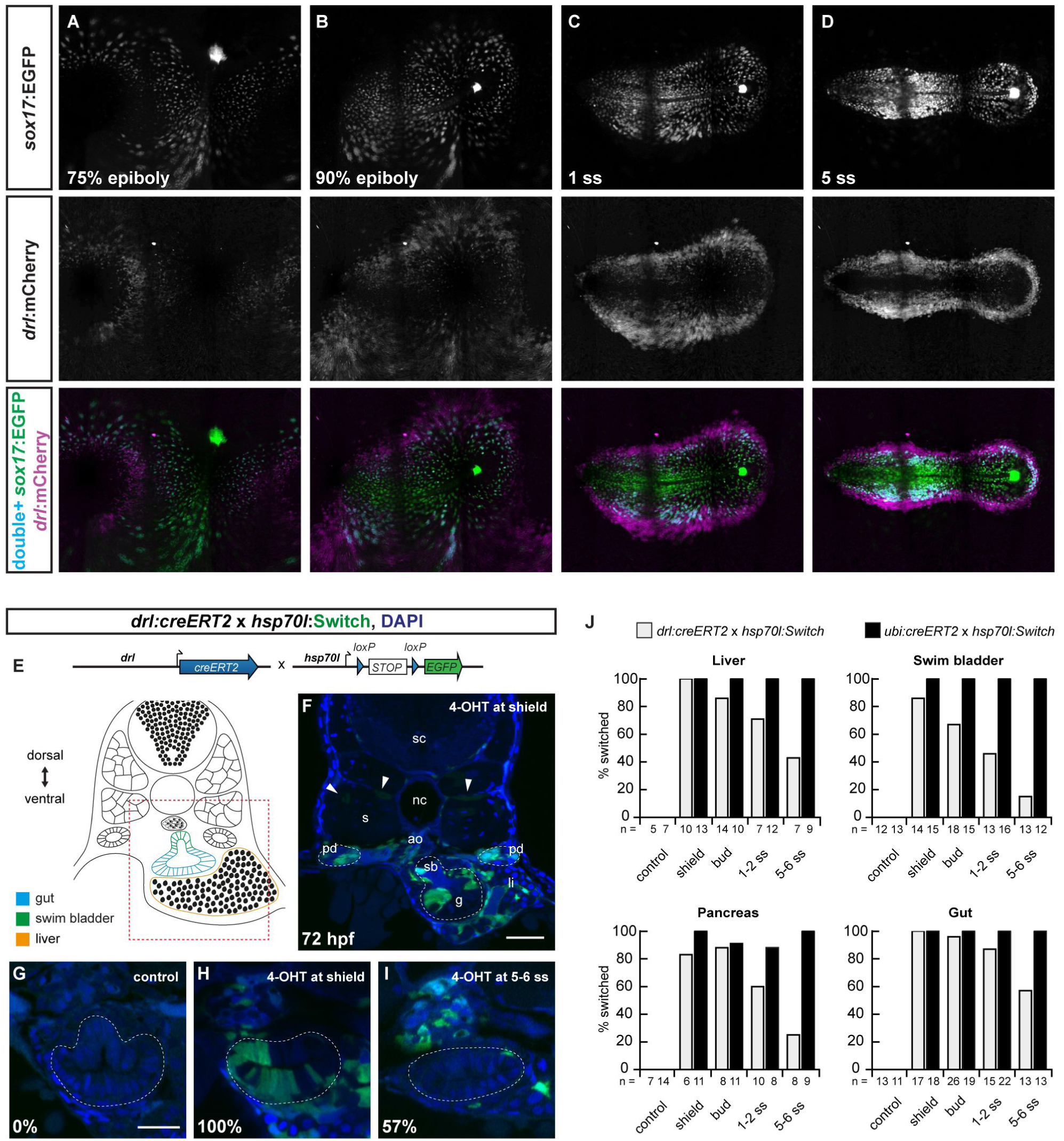
The early *drl* reporter-expressing cells comprise endoderm- and LPM-fated progenitors. (**A-D**) Single time point projections of *sox17:EGFP* marking endoderm progenitors (**A-D**) and *drl:mCherry* marking LPM progenitors from gastrulation until 5 ss; anterior to the left. Double-positive cells for EGFP and mCherry shown in blue (**A-D**). (**E**) Schematic of *drl:creERT2* to *hsp70l:Switch* cross for genetic lineage tracing and schematic of transverse section (trunk region) with endoderm- and LPM-derived organs at 3 dpf. (**F**) Representative 72 hpf transverse section of *drl* lineage-traced embryo 4-OHT-induced at shield stage; arrowheads depict rare trunk muscle labeling. (**G-I**) Transverse sections of *drl* lineage tracing at 72 hpf, control (**G**) versus 4-OHT-induced at shield stage (**H**) and 5-6 ss (**I**); note gradual loss of endoderm labeling (intestinal cells, dashed region). Numbers indicate percentage of embryos with intestinal lineage labeling. (**J**) Quantification of endoderm lineage labeling in representative organs following 4-OHT induction at indicated time points, comparing *drl:creERT2* versus ubiquitous *ubi:creERT2* control as reference for the *hsp70l:Switch* lineage reporter. Notochord (nc), somite (s), spinal cord (sc), pronephric duct (pd), liver (li), swim bladder (sb), gut (g), nuclei in blue (DAPI; **F-I**). Scale bar (**F**) 50 μm and (**G-I**) 25 μm.

In contrast, *sox17:creERT2* exclusively marked endoderm lineages (n = 6; Fig S1C), supporting data that *sox17* expression demarcates zebrafish endoderm progenitors downstream of the key endoderm regulator *sox32* (Alexander and Stainier, 1999). Remarkably, embryos that fail to form endoderm upon *sox32* perturbation still generated a type of *drl*-traced LPM that partitioned into its major recognizable descendants including heart, blood, endothelium, and pronephros (Fig S3). These results are not skewed by lineage-bias of *hsp70l:Switch* reporter sensitivity, as we observed well-distributed lineage labeling using the ubiquitous *ubi:creERT2* (Fig S1D).

Altogether, these data establish that during gastrulation the *drl*-marked LPM gradually refines from a ventral mesendoderm territory to a bilateral LPM domain as the sole mesodermal fate along the entire anterior-posterior axis of the embryo. The rare lineage labeling of somitic muscle by *drl:creERT2* (Fig 2F) (Gays et al., 2017; Mosimann et al., 2015) further underlines that, in zebrafish, the paraxial mesoderm and the LPM develop as distinct mesoderm lineages with only minimal overlap (Warga and Nüsslein-Volhard, 1999).

### A pan-LPM enhancer in the zebrafish *drl* locus

To identify *cis*-regulatory element(s) in the *drl* locus responsible for pan-LPM progenitor expression, we divided the 6.35 kb *drl* regulatory region into smaller fragments and assayed their activity using Tol2-based *EGFP* reporters in F0 and stable transgenics (Fig 3A). We found that the promoter-proximal region surrounding exon 1 recapitulated *drl* reporter expression in ALPM and PLPM from 5-7 ss onwards (*proximal drl*), while the upstream promoter region alone remained active in the posterior endothelial and blood precursors (−*1.02drl*) (Fig 3A-C). In addition, we identified a small (968 bp) region in the first intron (+*2.0drl*) that initially recapitulated early *drl* reporter expression in zebrafish embryos before fading between 5-10 ss (approx. 13 hpf) (Fig 3A,D). Genetic lineage tracing with +*2.0drl*:*creERT2* and *hsp70l:Switch* (Fig 3E) specifically labeled LPM-derived organs including heart, blood, endothelium, kidney, pectoral fin mesenchyme, and iSMCs, and additionally marked endoderm lineages when induced with 4-OHT at shield stage (Fig 3F-I). These results correlate well with our lineage tracing using full-length *drl:creERT2* initiated at the onset of gastrulation (Fig 2E-J) (Felker et al., 2018; Gays et al., 2017; Mosimann et al., 2015). Further, deletion of elements within the +*2.0drl* enhancer defined a minimal enhancer region of 432 bp (+*2.4drl*) that functioned as a pan-LPM enhancer, albeit with higher variability in stable transgenics (Fig S4). These regulatory analyses indicate that the entire *drl* expression pattern derives from distinct *cis*-regulatory elements that control *drl* expression in separable early mesendoderm/pan-LPM and later ALPM versus PLPM domains. The latter pattern is analogous to the hematopoietic lineages that arise during somitogenesis and are commonly marked with *drl* mRNA ISH (Davidson and Zon, 2004; Herbomel et al., 1999). This data implies that the +*2.0drl* enhancer contains the key regulatory modules that respond and integrate to early LPM-defining inputs.

**Figure 3:**
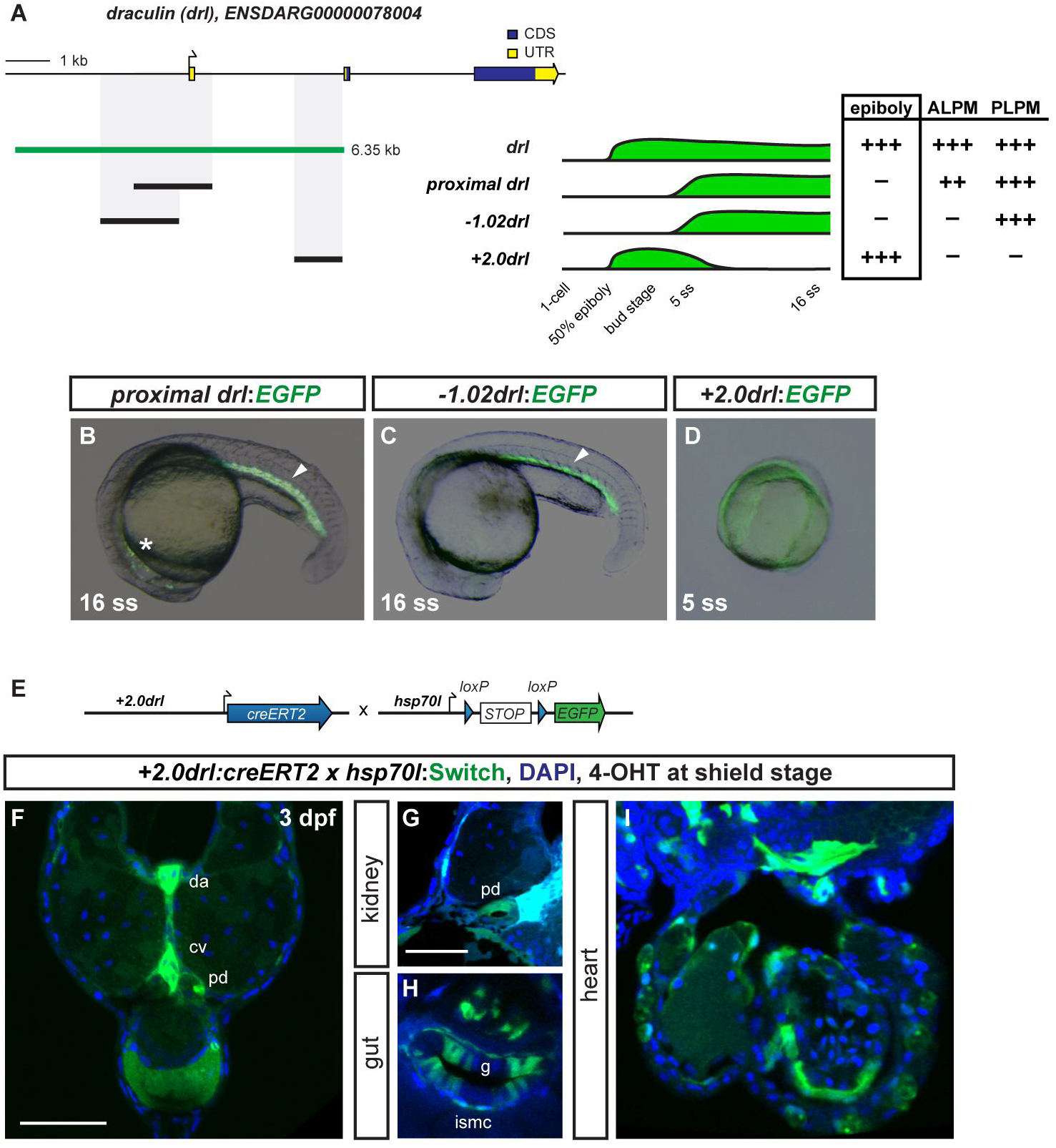
The 6.35 kb *drl cis*-regulatory region contains an early pan-LPM enhancer. (**A**) Schematic of the *drl* locus depicting the 6.35 kb *cis*-regulatory region (green), and smaller isolated candidate fragments *proximal drl* (region surrounding first exon), *−1.02drl* (upstream region only), and +*2.0drl* (distal first intron) with specific reporter activity. Time line and table indicates expression dynamics (50% epiboly to 16 ss) of stable transgenes for the individual regulatory elements and expression domains (pan-LPM early or somite-stage ALPM, PLPM) with absent expression (-) to strong expression (++). (**B-D**) Representative stable transgenic zebrafish embryos harboring EGFP reporters for *proximal drl, −1.02drl*, and +*2.0drl*; at 5 ss and 16 ss *proximal drl* and −*1.02drl* express in PLPM (arrowheads in **B,C**), *proximal drl* additionally in ALPM (asterisk in **B**); note pan-LPM activity of +*2.0drl* (**D**). (**E**) Schematic of +*2.0drl:creERT2* to *hsp70l:Switch* cross for genetic lineage tracing. (**F-I**) Transverse sections at 3 dpf of +*2.0drl:creERT2* lineage tracing after 4-OHT induction at shield stage results in specific labeling of LPM-derived tissue including heart (h), red blood cells (rbc), dorsal aorta (da) and cardinal vein (cv) endothelium, pronephric duct (pd), and intestinal smooth muscle cells (ismc). +*2.0drl:creERT2* also traces endoderm-derived tissue, shown for gut epithelium (g). Nuclei counterstained with DAPI (blue). Scale bars (**F**) 250 µm and (**G**) 50 µm.

### Combined EomesA, FoxH1, and MixL1 activity integrates ventral BMP activity into LPM formation

We next sought to investigate the upstream inputs that control the +*2.0drl* pan-LPM enhancer. BMP and Activin/Nodal ligands of the TGF-β superfamily trigger key pathways in early vertebrate axis determination and mesendoderm induction (Hill, 2018; Langdon and Mullins, 2011; Whitman, 2001). Both pathways interpret ligand gradients and signal through type I and type II receptors that result in cytosolic phosphorylation of Smad transcription factors (Fig 4A) (Massagué, 2012; Rogers and Müller, 2019). In zebrafish embryos during early gastrulation, BMP ligands are secreted from the ventral side, while Nodal ligands are expressed along the margin and the dorsal side (Hill, 2018; Langdon and Mullins, 2011). In line with the emergence of ventral LPM, endogenous *drl* expression was virtually absent in embryos maternally mutant for dominant-negative Smad5 (*MZsbn*), which lack BMP activity (Fig 4B,C) (Hild et al., 1999). Similarly, treatment with the BMP inhibitor Dorsomorphin resulted in pronounced decrease of endogenous *drl* expression (Fig 4E,F). We also found decreased *drl* expression in embryos with perturbed Nodal signaling: i) maternal-zygotic mutant embryos lacking the key Nodal co-receptor Crypto/Oep (*MZoep*) that cannot transmit Nodal signaling around the embryo margin (Gritsman et al., 1999; Schier et al., 1997), and ii) embryos treated with the Nodal signaling inhibitor SB-505124 (Hagos and Dougan, 2007) (Fig 4D,G,H). These results indicate that the ventral *drl* domain is sensitive to both BMP and Nodal input. Consistent with our lineage tracing results (Fig S3), embryos devoid of endoderm upon *sox32* knockdown still expressed endogenous *drl*, albeit with overall thinned-out expression and a marked decrease of dorsal *drl* activity (Fig 4I).

**Figure 4:**
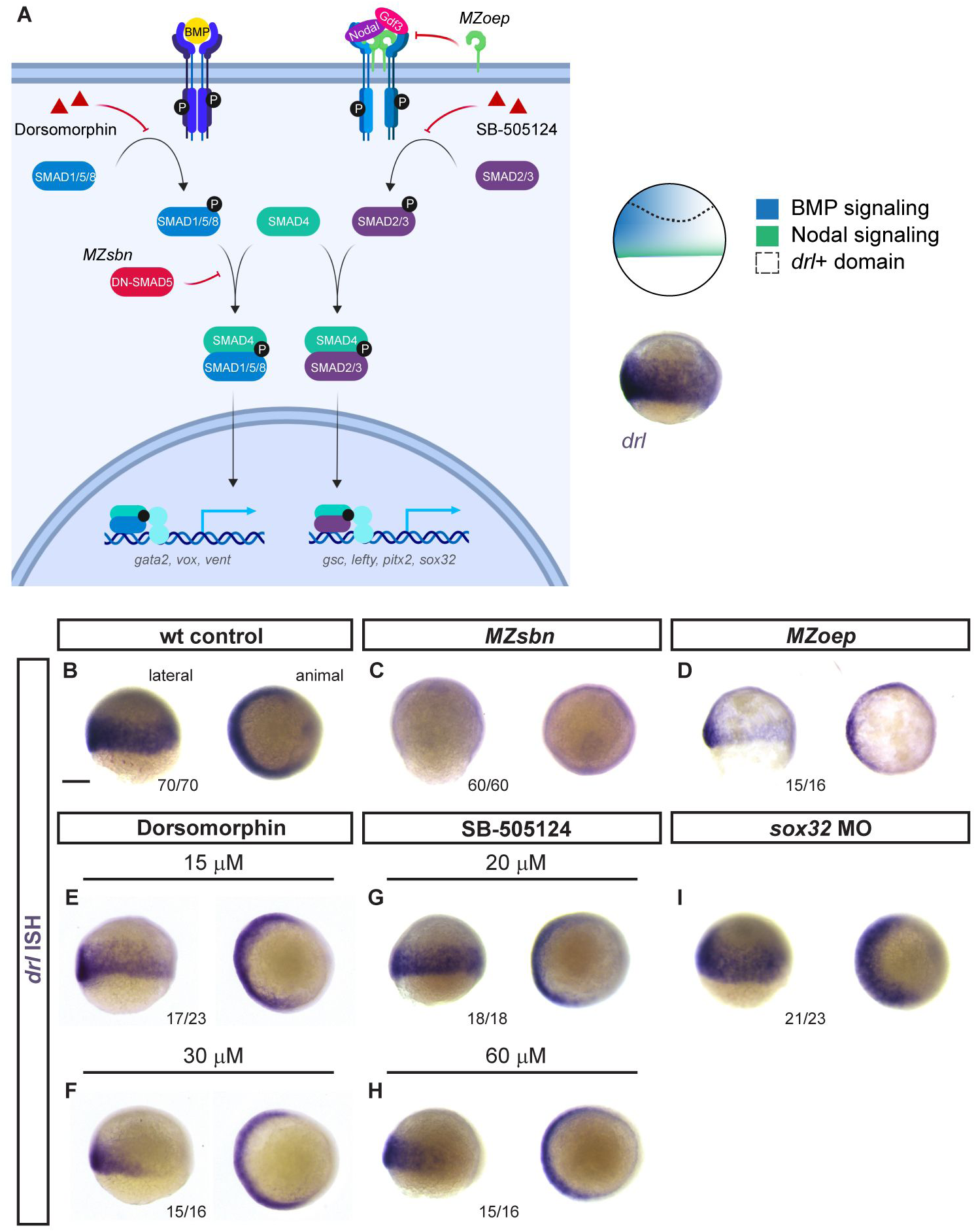
Early *drl*-expressing LPM progenitors respond to ventral BMP. (**A**) Schematic of the BMP and Nodal signaling pathways and the corresponding signaling territories in the gastrulation-stage zebrafish embryo, compared to endogenous *drl* expression marked by mRNA ISH. (**B-I**) mRNA ISH for endogenous *drl* expression during early gastrulation (shield stage to 75% epiboly), lateral on the left and animal view on the right. (**B-D**) Wildtype (*wt*) controls, BMP-perturbed (*MZsbn*, a dominant-negative Smad5 allele) and Nodal-perturbed (*MZoep*, mutant for the required Nodal co-receptor Oep) showing that BMP perturbation abolishes endogenous *drl* expression. (**E-H**) Chemical inhibition of BMP via Dorsomorphin (15 μM and 30 μM, **E,F**) decreases ventral *drl* expression, while chemical Nodal inhibition with SB-505124 (20 μM and 60 μM, **G,H**) decreases dorsal *drl* expression. (**I**) *sox32* morphants that fail to form endoderm also lose dorsal *drl* expression. Scale bar (**B**) 250 μm.

We mined published whole-embryo ChIP-seq data from zebrafish gastrulation stages (Dubrulle et al., 2015; Nelson et al., 2014; Nelson et al., 2017) and identified candidates for transcription factors binding to the +*2.0drl* enhancer. These include the T-box transcription factor EomesoderminA (EomesA), its interaction partner FoxH1, and BMP/Nodal-mediating Smads. Published evidence uncovered that they participate in control of mesendoderm genes (Charney et al., 2017; Chen et al., 1996; Germain et al., 2000; Slagle et al., 2011) and affect *drl* expression during early somitogenesis (Slagle et al., 2011) (Fig 5A). We found that mRNA injection- or *ubi* promoter-driven expression of constitutive-active forms of EomesA or FoxH1 strongly augmented and prolonged +*2.0drl* reporter and endogenous *drl* expression in their native LPM domain compared to controls (Fig 5B-E, Fig S5A-D). Ubiquitous expression of wildtype *eomesA* or *foxh1* mRNA was sufficient to increase endogenous *drl* expression (Fig 5F-H,). Addition of a constitutively-active Smad2 (Dick et al., 2000) to EomesA and FoxH1 resulted in dorsal widening of +*2.0drl* reporter expression pattern (Fig S5E,F).

**Figure 5:**
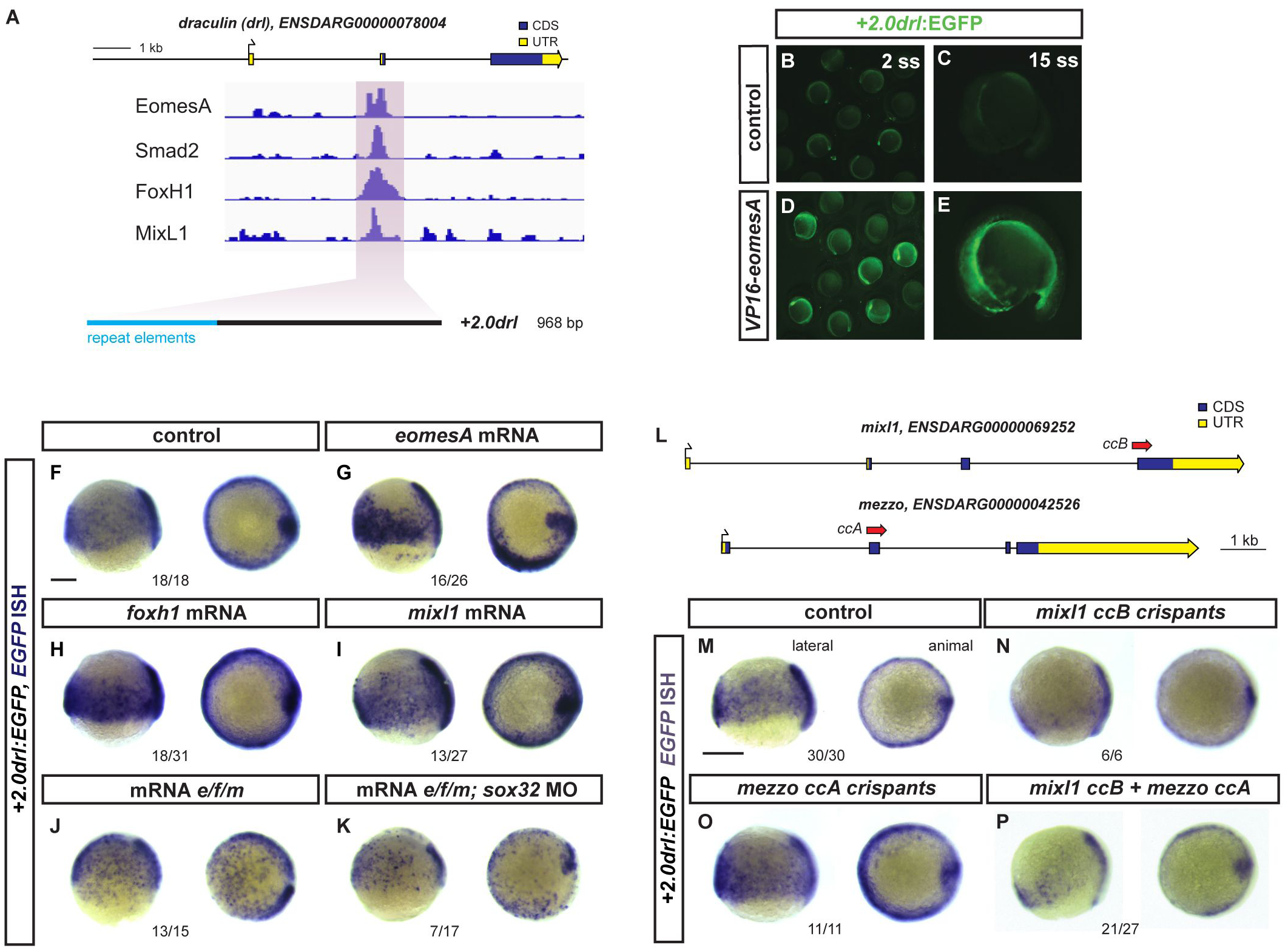
EomesA, FoxH1, and MixL1 together induce the +*2.0drl* enhancer. (**A**) ChIP-seq tracks for EomesA. FoxH1, MixL1, and Smad2 in the *drl* locus. See text for details. Bottom depicts the +*2.0drl* intronic enhancer and the smaller minimally specific region +*2.4drl* after removal of repeat sequences. (**B-E**) Constitutively-active VP16-EomesA boosts +*2.0drl:EGFP* reporter expression in its native territory. Compared to injection controls (**B,C**), microinjection of *VP16-eomesA* mRNA in +*2.0drl*:*EGFP* reporter transgenics enhances and prolongs EGFP expression in the native reporter expression domain (**D,E**). (**F-K**) Gastrulation-stage (shield to 75% epiboly) zebrafish embryos, lateral view left and animal view right, probed with *EGFP* ISH to detect expression of +*2.0drl:EGFP*. Compared to controls (**F**), embryos injected with mRNA encoding full-length *eomesA*, *foxh1*, or *mixl1* show enhanced +*2.0drl:EGFP* reporter activity (**G-I**). (**J**) Combining mRNAs encoding full-length *eomesA (e)*, *foxh1 (f)*, and *mixl1 (m)* (*e/f/m*) trigger ectopic reporter expression also in dorsal blastomeres (compare to native reporter expression pattern in **F**), an activity that also remained in embryos devoid of endoderm after *sox32* morpholino injection (**K**). (**L-P**) Mixl1 acts on the +*2.0drl* enhancer as analyzed in Cas9 RNP-mediated crispants. (**L**) Schematic representation of the *mixl1* and *mezzo* loci, with the individual sgRNAs for mutagenesis annotated (*cc* for *CRISPR cutting*, followed by sgRNA index). (**M-P**) mRNA *in situ* hybridization of *EGFP* expression in +*2.0drl*:*EGFP* embryos as crispant control (**M**), injected with Cas9 RNPs of (**N**) *mixl1 ccB*, (**O**) *mezzo ccA*, and (**P**) *mixl1 ccB* together with *mezzo ccA* sgRNA; the resulting mosaic *mixl1 ccB* crispants show diminished +*2.0drl* reporter expression at late gastrulation. Lateral and animal views as indicated. Scale bar in (**B**) 250 μm.

EomesA and FoxH1 are maternally contributed and are thus ubiquitously distributed during gastrulation (Bruce et al., 2003; Slagle et al., 2011), during which the LPM emerges in the ventral BMP and Smad activity domain (Fig 1,4). We therefore hypothesized that at least one additional, ventrally expressed transcription factor might be required for LPM formation. In published ChIP-seq data (Nelson et al., 2017), we identified the homeodomain protein MixL1 as a third possible transcription factor that acts together with EomesA and FoxH1 in controlling +*2.0drl* enhancer activity (Fig 5A). MixL1 is a downstream target of BMP and Nodal signaling implicated in controlling endoderm and mesoderm fates (Kunwar et al., 2003; Stainier et al., 1996) and it retains ventral expression in *MZoep* mutants (Slagle et al., 2011). Furthermore, MixL1 can form a complex with EomesA (Bjornson et al., 2005). Reminiscent of *eomesA* or *foxh1* mRNA injections (Fig 5G,H), microinjected *mixl1* mRNA also resulted in increased +*2.0drl* reporter expression within in the native LPM domain (Fig 5I).

Combining the triplet of wildtype mRNAs or Tol2-based DNA constructs encoding full-length EomesA, FoxH1, and MixL1 (shortened as *e/f/m*) led to ubiquitous +*2.0drl* reporter activation in embryos (Fig 5J). In *MZsbn*-mutant embryos without BMP signaling, *e/f/m* misexpression induced *drl* expression dorsally (Fig S5H). These observations suggest that in *e/f/m* overexpression conditions there is still a requirement for additional Smad activity, which in *MZsbn* embryos is only available in the dorsal Nodal-positive domain. Conversely, loss of Nodal signaling in *MZoep* mutants led to a ventral upregulation of *drl* expression upon mRNA-based *e/f/m* overexpression (Fig S5I). Combining native *e/f/m* in wildtype and *MZsbn* embryos devoid of endoderm following *sox32* knockdown also resulted in ubiquitous +*2.0drl* reporter activation (Fig 5K, Fig S5J). This suggests that most of the +*2.0drl* reporter-positive cells have an LPM identity. Mutating *mixl1* by CRISPR-Cas9 resulted in mosaic loss of +*2.0drl* reporter activity in F0 crispants (Fig 5L-N), while mutating the redundant *mixl1* paralog *mezzo* (Poulain and Lepage, 2002) alone or together with *mixl1* did not influence +*2.0drl* reporter activity (Fig 5L,O,P). This indicates that MixL1 is the predominant Mix paralog acting on the +*2.0drl* enhancer in zebrafish. Furthermore, CRISPR-Cas9-mediated mutagenesis of the +*2.0drl* enhancer in the region of predicted FoxH1 and MixL1 sites in the context of the full-length *drl:EGFP* transgene resulted in specific perturbation of early LPM reporter expression, without affecting the later ALPM and PLPM patterns (Fig S6).

EomesA, FoxH1, and MixL1 misexpression also induced expression of *tmem88a*, which is highly enriched in the early native LPM (Mosimann et al., 2015) (Fig S7A,B). In contrast, the expression domains of other early expressed mesodermal genes either showed a slight broadening of their native domains or appeared generally unaffected (Fig S7). This is further illustrated by the lack of changes in *hand2* expression, which normally initiates in the lateral-most LPM after gastrulation (Fig S7K-M). Together, these data suggest a regulatory model in zebrafish whereby the combination of EomesA, FoxH1, and MixL1 potentiates ventral Smad-relayed BMP signals to demarcate a mesendoderm territory that becomes prospective LPM.

### The *drl* pan-LPM enhancer reads out an LPM program across chordates

We next explored whether the +*2.0drl* enhancer could read out a putative pan-LPM program in diverse chordate species. First, we tested several previously characterized enhancers with activity in the posterior LPM of mice: G*ata4* (Rojas et al., 2005), *Bmp4* (Chandler et al., 2009), and *HoxB6* (Becker et al., 1996). Reporter transgenes based on mouse *Gata4* and *Bmp4* showed restricted activity in the outward migrating endothelial/blood progenitors and in the PLPM when electroporated into the primitive streak of *ex ovo*-cultured chicken embryos after the onset of gastrulation (HH3+/4) (Fig S8B,C). The mouse *HoxB6* LPM enhancer showed no regulatory activity in this assay (Fig S8D). In contrast, when microinjected into zebrafish embryos, reporters based on these three mouse enhancers all resulted in expression mainly in the notochord without specific LPM activity (Fig S8E-H). This indicates that while some of the previously isolated LPM enhancers of mouse expressed faithfully in the PLPM of chick embryos, their activity did not recapitulate an LPM pattern in zebrafish. These results suggest that these mammalian LPM enhancers may have specialized during amniote evolution.

Electroporation of the zebrafish +*2.0drl* reporter into the primitive streak of HH3+/4 chicken embryos resulted in reporter activity specifically in the forming LPM: depending on the exact stage and region of electroporation, we observed specific reporter activity in several LPM territories. Most frequently observed expression patterns included medial and posterior LPM domains (Fig 6A-C, Fig S9), and we also observed ALPM reaching the head fold in individual embryos (Fig S9). These observations suggest that a basic upstream program underlying LPM formation, as read out by the +*2.0drl* reporter, continues to function in birds as representative amniotes.

**Figure 6:**
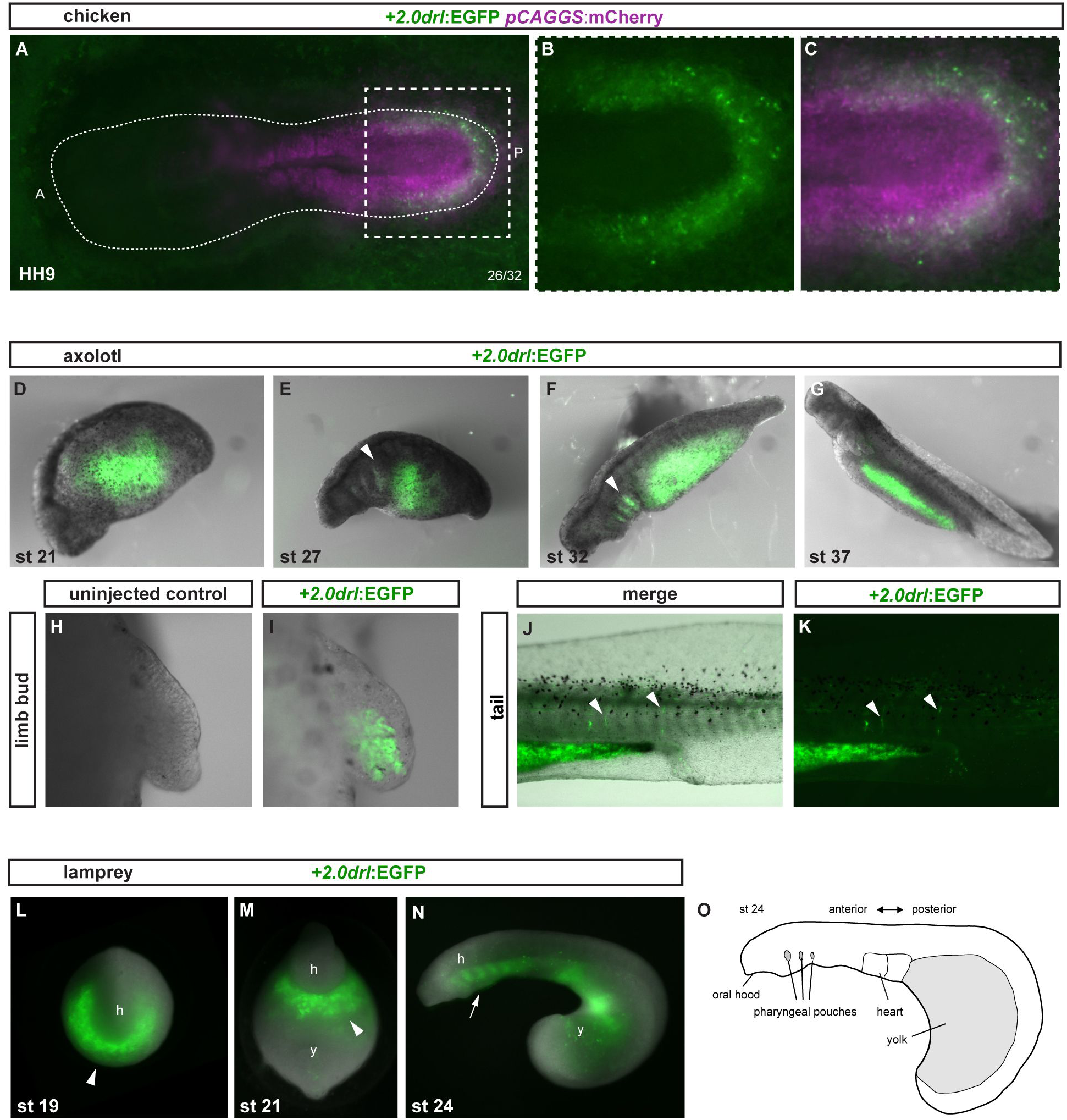
The zebrafish +*2.0 drl cis*-regulatory element reads out a LPM program across vertebrates. (**A-C**) HH9 *ex ovo*-cultured chicken embryo electroporated at HH3+/4 with +*2.0drl*:EGFP (green, **A-C**) and ubiquitous *pCAGGS:mCherry* control (magenta, **A,C**), showing specific +*2.0drl* reporter expression in the electroporated LPM. The dashed line depicts the outline of the chicken embryo, anterior to the left; boxed region (**A**) depicts magnified area shown for single and merged channels (**B,C**).(**D-G**) Expression of +2.0*drl*:EGFP at the indicated embryonic stages. Note EGFP expression in the lateral portion of the embryo, future gut region, and pharyngeal arches (arrowheads in **E,F**). (**H,I**) EGFP expression in mesenchymal cells of the developing limb bud, indicative of LPM origin. (**J,K**) EGFP fluorescence in stage 43 larvae in the gut lining and blood vessels (arrowhead). Expression is occasionally found in a small fraction of muscle fibers. (**L-O**) Transient transgenic lamprey embryos (*Petromyzon marinus*) with +*2.0drl*:EGFP expression in the anterior mesendoderm (arrowheads) and overlying the yolk at neurula stages (st 19-21) (**L,M**), and in the developing pharynx (arrow) during head protrusion (st 23-24) (**N**); views anterior (**L**), ventral (**M**), lateral (**N**), head (h) and yolk (y). (**O**) Schematic depiction of st23 lamprey embryo to outline key features.

We then tested whether +*2.0drl* is responding to LPM-inducing cues in other tetrapods. Axolotl embryos microinjected with the +2.0*drl* EGFP reporter marked putative endodermal and LPM territories beginning from early somite-stages (stage 21), additionally marking the pharyngeal regions at tailbud stages (st 27 and 32 shown in Fig 6D-G). Transversal sections of stage 32 embryos confirmed the presence of EGFP-positive cells in the endoderm (asterisk) and lateral mesendoderm (Fig S9G-J). Interestingly, EGFP fluorescence was present throughout axolotl development and could be readily detected in the gut, as well as LPM-derived tissues including the limb bud (Fig 6H-I), blood vessels (Fig 6J-K), and gut lining (Fig S9K-N). These results support the notion that the +2.0*drl* enhancer also interprets an LPM program active in amphibians.

Next, we asked if the +*2.0drl* enhancer reads out a pan-LPM program in more distantly related vertebrates. Lampreys are jawless vertebrates (cyclostomata) that can provide unique insights into vertebrate evolution due to the early divergence of their lineage from jawed vertebrates (Shimeld and Donoghue, 2012). Microinjection of the +*2.0drl* EGFP-based reporter into sea lamprey embryos (*Petromyzon marinus*) consistently resulted in robust EGFP expression in the lateral mesendoderm starting during neurulation (st 18-21) (n = 145/231) (Fig 6L,M), as well as in the developing pharynx at st 22-24 (Fig 6N). Transverse embryo sections revealed that this early expression domain includes the anterior-most, LPM-linked expression of lamprey *pmHandA* (Onimaru et al., 2011), with the later pharyngeal expression of EGFP being restricted to the endoderm and mesoderm (Fig S10A-K). We conclude that the +*2.0drl* enhancer is capable of integrating regulatory outputs from an upstream LPM program that remains conserved across vertebrates.

Next, we asked if the +*2.0drl* enhancer also responds to upstream activity in LPM-linked cell fates dating back to the chordate radiation (Fig 7A). We first electroporated the +*2.0drl* reporter into embryos of the tunicate *Ciona robusta*, a chordate species belonging to a sister clade of vertebrates (Fig 7A). While missing the full complement of LPM-derived organ systems found in vertebrates, the LPM is echoed in the cardiopharyngeal progenitors forming in *Ciona* embryos (Diogo et al., 2015; Kaplan et al., 2015). We detected +*2.0drl:EGFP* reporter activity in emerging cardiac and pharyngeal muscle lineages at *Ciona* larval stage (st 26; 18 hpf (Hotta et al., 2007)) (Fig 7B-D): we observed +*2.0drl*:EGFP reporter activity in the atrial siphon muscle precursors (ASMPs) and in both first and second heart precursors (FHPs and SHPs). This was confirmed by co-localization of *Mesp:H2B-mCherry* expression that labels the cardiopharyngeal cell lineage (Davidson et al., 2005) (n =15/92; Figure S10L-M). In agreement with the *drl*-based LPM lineage tracing in zebrafish, we found minimal to no overlap with paraxial mesoderm progenitors, and rarely observed EGFP expression in the anterior tail muscles (ATMs) of the *Ciona* larval tail (Fig 7B-D). These results indicate that the zebrafish +2.0drl enhancer responds to regulatory input in the emerging multipotent cardiopharyngeal progenitors in *Ciona*.

**Figure 7:**
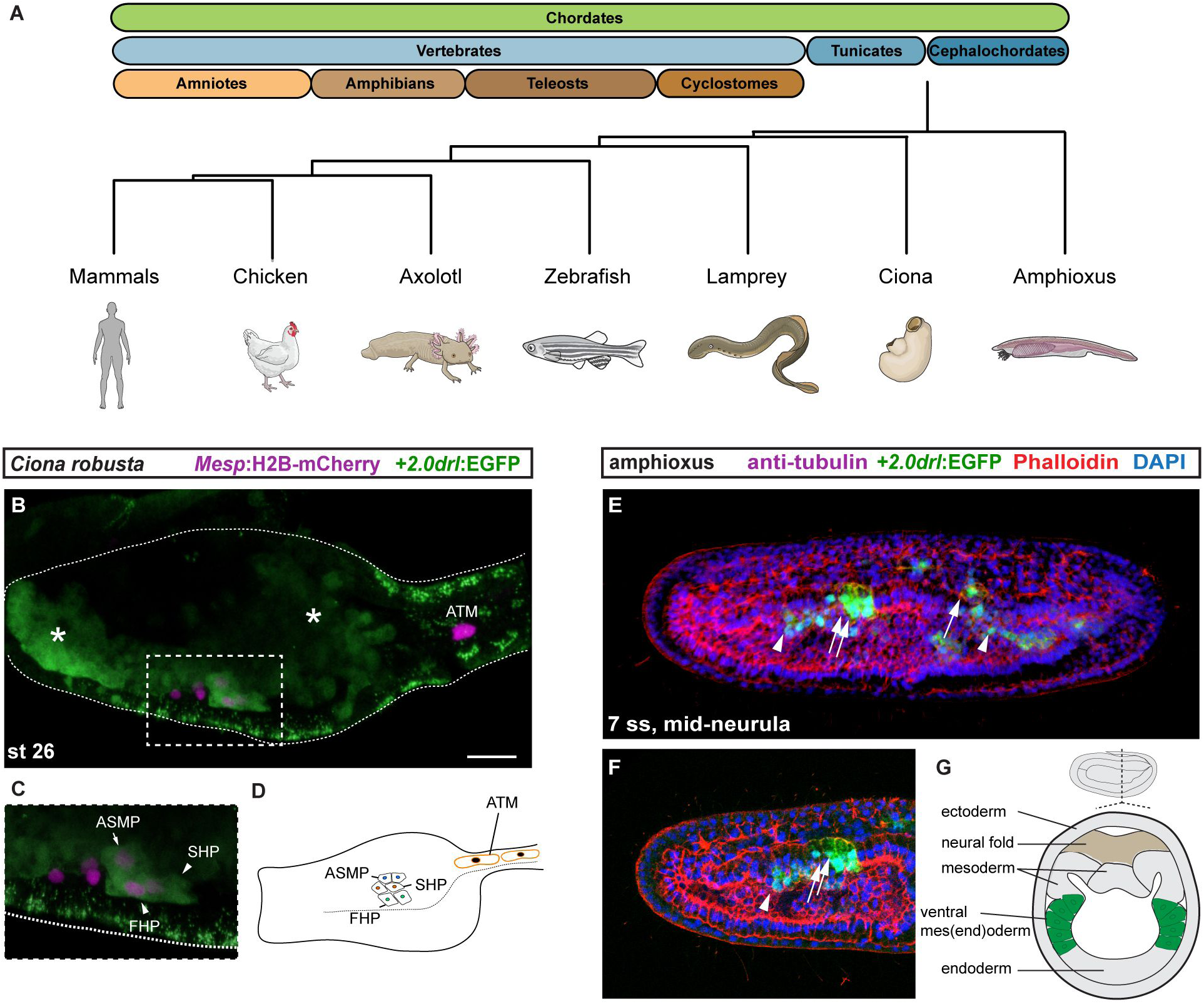
The zebrafish +*2.0drl cis*-regulatory element reads out an LPM program in tunicates and cephalochordates. (**A**) Phylogeny of chordates depicting the species used in this study. (**B-D**) *Ciona robusta* embryo at 18 hpf (st 26), electroporated with +*2.0drl*:EGFP (green) and with *Mesp*:H2B-mCherry (magenta) to track the B7.5 cardiopharyngeal cell lineage. (**B,C**), representative larva shown with boxed region (B) magnified for detail (**C**); dashed line indicates midline, anterior to the left, schematic larva depicted in (**D**). +*2.0drl*-driven EGFP partly overlaps with B7.5 derivatives including atrial siphon muscles precursors (ASMP) (white arrows) and both first and second heart precursors (FHPs and SHPs) (white arrowheads). EGFP is also detected in mesenchymal lineages (white asterisks). (**E-G**) Mid-neurula stage (7 ss) amphioxus embryo, confocal Z-stack anterior to the left and dorsal to the top; embryo injected with +*2.0drl:EGFP* (green), counterstained with Phalloidin (red), DAPI (blue), lateral view as 3D-rendering (**E**) and Z-stack sagittal section (**F**). +*2.0drl:EGFP* showing specific reporter activity in lateral endoderm (arrowhead), ventral half of somites (arrows), and elongating somites (double arrows). (**G**) Schematic depiction of amphioxus embryo; lateral view on top, dotted line represents transverse section depicted below with green depicting domain of +*2.0drl:EGFP* expression. Scale bar (**B**) 25 μm.

Lastly, we examined the cephalochordate amphioxus (*Branchiostoma lanceolatum*), which belongs to the most basally divergent lineage of chordates (Bertrand et al., 2011). In amphioxus, the LPM forms from a continuous sheet of cells that encompass the dorsally emerging somites, the LPM, and the ventral-most forming endoderm (Bailey and Miller, 1921; Holland, 2018; Holland et al., 2003; Kozmik et al., 2001; Onimaru et al., 2011). At mid-neurula stage (approximately equivalent to early somitogenesis in zebrafish), the ventral wall of the somites evaginates as nascent ventral mesoderm, which by late neurula stage fuses at the midline under the gut endoderm (Holland et al., 2003; Kozmik et al., 2001; Onimaru et al., 2011). Indeed, the amphioxus orthologs of LPM-expressed transcription factors including *Hand*, *Csx*, *Vent1*, and *AmphiNk2-tin* are expressed in the ventral half of the somite territory at mid-neurula stage (Holland et al., 2003; Kozmik et al., 2001; Pascual-Anaya et al., 2013). We observed that injection of +*2.0drl*-based reporters into amphioxus embryos showed specific reporter activity in the ventral half of somites and in the elongating somites at mid-neurula stage (6/7 ss) (n = 61/65) (Fig 7E-G, Fig S10N,O). At early larvae stage the activity of +*2.0drl*-based reporter is present in the pharyngeal region (Fig S10P) where LPM is located (Holland, 2018). Hence, also in amphioxus as cephalochordate, the +*2.0drl* enhancer reads out the positional input active in the LPM-corresponding territory during development.

Taken together, these observations establish that the intronic +*2.0drl* enhancer reads out a position-dependent LPM program that remains active in tunicates, cyclostomes, teleosts, amphibians, and amniotes, and thus across all tested chordates.

## Discussion

The dynamic nature of the LPM has made it challenging to monitor precisely its emergence and morphogenesis and has hindered comparative studies of its properties during chordate evolution. Here, we show that an enhancer from the zebrafish *drl* locus (+*2.0drl*) reads out an LPM-demarcating transcriptional activity in six chordate species, ranging from cephalochordates to amniotes. This suggests that the LPM-underlying transcriptional program is of ancient evolutionary origin. Characterization of the properties of this enhancer in zebrafish revealed that the transcription factors Eomes, FoxH1, and MixL1 are sufficient to trigger this basic LPM program. These observations in zebrafish suggest a regulatory model whereby, among their roles in mesendoderm regulation, Eomes, FoxH1, and Mixl1 cooperate in inducing LPM together with position-dependent Smad activity. These factors have been individually implicated in mesendoderm development in several vertebrates (Arnold et al., 2008; Bjornson et al., 2005; Germain et al., 2000; Nelson et al., 2014; Slagle et al., 2011; Zhang et al., 2009), in diverse LPM-associated contexts such as in blood formation (Mead et al., 1996) and in reprogramming towards cardiac and renal fates (Costello et al., 2011; Pfeiffer et al., 2018; Takasato and Little, 2015; Takasato et al., 2014). Based on our series of comparative and mechanistic studies, we postulate that LPM-like origins in ancestral chordates already featured the basic molecular building blocks that enabled the increasing specialization of the LPM into its sophisticated descendant cell fates observed in vertebrates.

### The LPM as early established, specialized mesoderm

In vertebrates, the LPM is readily detectable after gastrulation through its position lateral to the forming somites and by several progenitor markers for hemangioblast, renal, and smooth muscle fates (Jin et al., 2006; McDole et al., 2018; Picker et al., 2002; Yin et al., 2010; Zhu et al., 2005). Genetic tracking of some aspects of the LPM has been achieved previously by various means in different models. In mouse, transgenic strains based on *HoxB6*, *Prrx1*, *Bmp4*, *Hand1/2*, *Gata2*, and *FoxF1* enable labeling of the LPM post-gastrulation (Becker et al., 1996; Chandler et al., 2009; Firulli et al., 1998; Martin and Olson, 2000; Ormestad et al., 2004; Osterwalder et al., 2014; Rojas et al., 2005). Recent work in chick used electroporation of ubiquitous reporters specifically at the position of emerging LPM progenitors to chart the forming limb and interlimb domains (Moreau et al., 2019). In zebrafish, we had previously found and applied the *cis*-regulatory region of the *drl* gene to genetically track LPM emergence during both gastrulation and early somitogenesis (Felker et al., 2018; Gays et al., 2017; Henninger et al., 2017; Mosimann et al., 2015). While *drl* encodes a putative zinc-finger protein of unknown function (Herbomel et al., 1999) without a clear ortholog outside of zebrafish, the early LPM-confined expression mediated by its +*2.0drl* enhancer provides a unique tool to investigate LPM origins across chordates. Using *in toto* live imaging of *drl*-based reporters together with lineage-restricted reporter transgenics, we charted LPM formation in zebrafish as a continuous process building around the entire circumference of the forming embryo (Fig 1, Movie S1). This mode of progenitor formation is distinct from paraxial and axial mesoderm, which both form by progressive extension over time (Alev et al., 2013; Gurdon, 1995).

Our reporter and lineage tracing data documents a close developmental relationship between endoderm and the LPM. Expression of the key vertebrate endoderm gene *sox17* demarcates prospective endoderm progenitors also in zebrafish (Alexander and Stainier, 1999; Sakaguchi et al., 2006), as evident by the exclusive endoderm lineage-labeling with *sox17:creERT2* (Fig S1C) (Hockman et al., 2017). We detected *sox17*/*drl* reporter double-positive cells throughout gastrulation that begin to separate from each other from tailbud stage (Fig 2A-D). Cre/*lox*-based lineage tracing confirmed that *drl:creERT2*-expressing cells at shield stage contribute to all endoderm-derived organs, while their mesodermal contribution was exclusively to the LPM (Fig 2F). The endoderm contribution gradually decreased from mid-gastrulation to early somitogenesis in an anterior-to-posterior fashion, while LPM derivatives remained labeled throughout (Fig 2E-J, Fig S1, S2). These findings indicate that *drl* labels a mesendoderm population that becomes progressively dedicated towards an LPM fate in a spatio-temporal manner.

Despite this close developmental relationship, *drl*-labeled LPM progenitors do not seem to require endoderm for their initial morphogenesis. While LPM midline migration is perturbed in *sox32* (*casanova*) mutants or morphants devoid of endoderm progenitors, such embryos still form bilateral, contracting hearts (Alexander et al., 1999; Dickmeis et al., 2001; Kikuchi et al., 2001) and maintain kidney and iSMC progenitor markers (Chou et al., 2016; Reichenbach et al., 2008). Our LPM lineage tracing confirmed that the LPM stripes still form even without endoderm and it documented how they migrate in the trunk to the midline and develop into structures resembling pronephros, iSMC-like structures, endothelium, red blood cells, and pectoral fin mesenchyme (Fig S3). These data imply that, despite close or even joint origin, the *sox17*-positive endoderm progenitors have minimal influence on initial LPM fate determination and LPM morphogenesis. Our imaging and lineage tracing data further indicates minimal overlap between early paraxial mesoderm progenitors and LPM progenitors, as evident in the rare occurrence of somatic muscle labeling by *drl*-expressing precursors (Fig 2F). While there is considerable heterogeneity of cell fate domains among the post-gastrulation LPM (Fig 1), our findings collectively suggest that the LPM initially emerges as a field of cells endowed with common properties.

### A transcription factor code for mesendoderm-to-LPM formation

Guiding the therapeutically relevant differentiation of cultured embryonic or induced pluripotent stem (iPS) cells towards cardiovascular, hematopoietic, or renal cell fates remains challenging (De Los Angeles and Daley, 2013; Lee et al., 2017; Slukvin II, 2013; Song et al., 2012; Takasato et al., 2014). Initial differentiation leads to broadly defined mesodermal progenitors that, depending on the protocol, show a preference to early versus late primitive streak regions, mimicking the anterior-to-posterior progression of vertebrate body axis formation (Mendjan et al., 2014). Other protocols combined expression of transcription factor combinations to drive direct differentiation into specific cell fates, such as achieved for cardiomyocytes or kidney cells (Ieda et al., 2010; Song et al., 2012; Takasato and Little, 2015; Takeuchi and Bruneau, 2009). Nonetheless, directed differentiation of uncommitted cells into correct LPM progenitor states, such as mesendoderm or mesoderm, would be highly desirable to achieve increased efficiency in cardiomyocyte, blood, or kidney reprogramming (Chia et al., 2019; Lee et al., 2017; Murry and Keller, 2008). In this regard, our functional analyses in zebrafish showed that Eomes, FoxH1, and MixL1 together with BMP-induced Smads are able to drive cells towards an LPM program.

Eomes and FoxH1 cooperate in controlling BMP/Nodal target genes together with Smads(Arnold et al., 2008; Bjornson et al., 2005; Miyazono et al., 2018; Slagle et al., 2011). Eomes, FoxH1, and Mixl1 have been implicated separately or in pairwise combinations in mesendoderm development (Chen et al., 1996; Du et al., 2012; Henry and Melton, 1998; Kikuchi et al., 2000; Slagle et al., 2011; Xu et al., 2014). In particular, Eomes and MixL1 have been found to form a tripartite complex with Gata5 in early endoderm determination (Bjornson et al., 2005). Our findings reveal that the combination of Eomes, FoxH1, and Mixl1 modulates mesendodermal target genes required for progression towards LPM formation. The requirement for the combined action of all three factors becomes apparent when testing each factor individually, as there was only a marginal increase in +*2.0drl* expression. In contrast, the combination of all three factors was sufficient to ubiquitously induce the +*2.0drl* pan-LPM reporter (Fig 5K) and *tmem88a*, which is an early LPM marker gene (Fig S7A,B) (Mosimann et al., 2015). The three factors do however not merely boost mesendoderm fate *per se*, as demonstrated in embryos devoid of endodermal progenitors following *sox32* perturbation (Fig 5L). The dependency of early *drl* expression on BMP, an less so on Nodal based on comparing genetic mutants, is further in line with the classic definition of a ventral emergence of the LPM (Gurdon, 1995). From this data, we propose the following model: maternal Eomes and FoxH1 cooperate with BMP- and Nodal-triggered Smads to prime mesendodermal target genes. Ventral induction of MixL1 provides an instructive signal that cooperates with the previous permissive mesendoderm state to trigger an LPM fate in ventral BMP-receiving blastomeres (Fig 4E-I). These findings provide a developmental framework for the contribution of Eomes, FoxH1, and MixL1 in programming of naïve pluripotent stem cells into cardiovascular and renal lineages by generating the correct mesendodermal precursor lineage. In-depth analysis of genomic targets of the three transcription factors is warranted to i) establish how this program conveys key LPM properties to uncommitted progenitor cells for their differentiation, and ii) if or which orthologs of these T-box, Forkhead, and Homeobox factors drive LPM progenitor formation across chordates.

### An ancient LPM program dating back to chordate ancestors

The evolutionary origin of the LPM has remained unaddressed. In part, the discussion of evolutionary origins of key features in the vertebrate body plan is tangled by the deduction of ancestral versus derived features without an existing common chordate ancestor (Technau and Scholz, 2003). Jawed vertebrate species share thousands of conserved non-coding regulatory regions (McEwen et al., 2009) and a greatly reduced number can be traced to jawless vertebrates like lamprey(Parker et al., 2011). Nonetheless, while some of the previously isolated LPM enhancers of mouse *Gata4*, *Bmp4*, and *Hoxb6* (Becker et al., 1996; Chandler et al., 2009; Rojas et al., 2005) expressed faithfully in the PLPM of chick embryos, their activity did not recapitulate an LPM pattern in zebrafish (Fig S8). These observations suggest that these mammalian LPM enhancers have specialized during amniote evolution. In contrast, our cross-species regulatory analyses of the +*2.0drl* enhancer uncovered a remarkable degree of regulatory conservation. While cryptic reporter activity in cross-species assays can bias results, we established that the zebrafish +*2.0drl* enhancer drove specific fluorescent reporter expression in LPM or LPM-related structures in six analyzed chordates: chick, axolotl, zebrafish, lamprey, *Ciona*, and amphioxus (Fig 6,7). Of note, in *Ciona*, the +*2.0drl:EGFP* reporter also resulted in EGFP activity in cells of the developing mesenchyme; if this reporter activity is specific considering the lineage origin of this cell type (Beh et al., 2007) or ectopic activity of the used transgene plasmid as previously reported (Stolfi and Christiaen, 2012), remains to be determined. Nonetheless, the +*2.0drl* enhancer provides a unique tool to investigate the upstream regulatory networks and the emergence of LPM structures across chordate development. This remarkable conservation of upstream reporter inputs further suggests that +*2.0drl* enhancer activity uncovers a deeply rooted LPM-inducing program, dating back to the last shared chordate ancestor.

Our findings further provide a genetic approach for investigating an LPM-delineating program that sets this mesodermal lineage apart from axial and paraxial mesoderm progenitors. The LPM program responds to ancient upstream regulatory inputs that defines the LPM from its early developmental origins across chordates. The activity of the +*2.0drl* reporter in the prospective LPM of amphioxus is particularly striking (Fig 7), as cephalochordates form few and only rudimentary equivalents of the vertebrate LPM-derived organ systems. The LPM in amphioxus forms as a non-segmented mesodermal sheet that is continuous with the ventral prospective endoderm and the more dorsally folding somites (Fig 7) (Bailey and Miller, 1921; Holland, 2018; Holland et al., 2003; Kozmik et al., 2001). This configuration makes it tempting to speculate that the LPM evolved from mesenchymal mesendoderm that did not integrate into the definitive endoderm or into the paraxial somites, providing ample material for diversification over deep time.

## Acknowledgements

We thank Seraina Bötschi, Lukas Obernosterer, and Vesna Barros for technical and husbandry support; the lab of Dr. Stephan Neuhauss for zebrafish husbandry support; the labs of Dr. Esther Stöckli and Dr. Jerome Gros for chicken experimentation support; the ZMB at UZH for imaging support; Dr. Fiona Wardle for input and support on the ChIP-seq panel in Figure 5A; the lab of Dr. Magdalini Polymenidou for vibratome access; Karolína Ditrychová for cloning the *pKD001* construct; Dr. Ashley Bruce, Dr. Rebecca Burdine, and Dr. Michael Tsang for sharing transcription factor constructs; Mark Miller for animal illustrations; the Stowers Institute histology facility for assistance with lamprey embryo sectioning; Dr. Hans-Henning Epperlein for discussions on salamander embryology; and all members of the Mosimann lab for constructive input. This work has been supported by a Swiss National Science Foundation (SNSF) professorship [PP00P3_139093] and SNSF R’Equip grant 150838 (Lightsheet Fluorescence Microscopy), a Marie Curie Career Integration Grant from the European Commission [CIG PCIG14-GA-2013-631984], the Canton of Zürich, the UZH Foundation for Research in Science and the Humanities, the Swiss Heart Foundation, and the ZUNIV FAN/UZH Alumni to C.M; a UZH CanDoc to C.H.; EuFishBioMed and Company of Biologists travel fellowships to K.D.P.; the Stowers Institute (grant #1001) to H.J.P. and R.K.; NIH/NHLBI R01 award HL108643, trans-Atlantic network of excellence award 15CVD01 from the Leducq Foundation to L.C.; a long-term fellowship ALTF 1608-2014 from EMBO to C.R; Alexander von Humboldt fellowship to A.C. and DFG Research Center (DFG FZ 111) and Cluster of Excellence (DFG EXC 168) funds to M.H.Y.; Czech Science Foundation 17-15374S to Z.K.

## Author Contributions

K.D.P., C.H., S.N., E.C.B., S.B., C.M. designed, performed, and interpreted zebrafish experiments; S.N., E.C., A.B. established and performed chicken experiments; K.D.P. performed lightsheet imaging with technical and equipment support by G.S., J.H.; K.W.R., P.M. provided and generated mutants and maternal-zygotic mutant zebrafish; H.J.P., M.B., R.K. designed, performed, and interpreted lamprey experiments; A.C., D.K., M.H.Y. designed, performed, and interpreted axolotl experiments; C.R., L.C. designed, performed, and interpreted *Ciona* experiments; I.K., Z.K. designed, performed, and interpreted amphioxus experiments; A.C., D.K., and M.H.J. designed and performed axolotl experiments; K.D.P., C.H., S.N., and C.M. assembled and wrote the manuscript with contributions from all co-authors.

## Materials and Methods

### Animal experiments and husbandry

Zebrafish and chick experiments were carried out in accordance with the recommendations of the national authorities of Switzerland (Animal Protection Ordinance). The protocols and the experiments were approved by the cantonal veterinary office of the Canton Zurich (Kantonales Veterinäramt, permit no. 150). Zebrafish care and all experimental procedures were carried out in accordance with the European Communities Council Directive (86/609/EEC), according to which all embryo experiments performed before 120 hours post fertilization are not considered animal experimentation and do not require ethics approval. Adult zebrafish for breeding were kept and handled according to animal care regulation of the Kantonales Veterinäramt Zürich (TV4209). All zebrafish (*Danio rerio*) were raised, kept, and handled as described (Westerfield, 2007). White mutant (d/d) axolotls (*Ambystoma mexicanum*) were obtained from the axolotl facility at the TUD-CRTD Center for Regenerative Therapies Dresden, Germany.

Lamprey studies were conducted in accordance with the Guide for the Care and Use of Laboratory Animals of the National Institutes of Health, and protocols were approved by the Institutional Animal Care and Use Committees of the California Institute of Technology (Protocol # 1436-11).

### Transgenic constructs and transgenic zebrafish lines

The upstream *cis*-regulatory elements of the zebrafish *drl* gene (*ENSDARG00000078004*; *ZDB-GENE-991213-3*) were amplified from zebrafish wildtype genomic DNA (Extended Data Table 1; regulatory elements) and TOPO-cloned into the *pENTR™ 5’-TOPO® TA* Cloning® plasmid (Invitrogen) according to the manufacturer’s instructions.

Subsequent cloning reactions were performed with the Multisite Gateway system with LR Clonase II Plus (Life Technologies) according to the manufacturer’s instructions. −*1.0drl:EGFP (pDestTol2pA2_-1.0drl:EGFP)* and [*proximal*]*drl:EGFP (pDestTol2pA2_*[*proximal*]*drl:EGFP)* were assembled from pENTR/5’_−*1.0drl* or pENTR/5’_[*proximal*]*drl* together with Tol2kit vectors *#383* (*pME-EGFP*), *#302* (*p3E_SV40polyA*), and *#394* (*pDestTol2A2*) as backbone. +*2.0drl:EGFP (pDestTol2pA2_+2.0drl:EGFP)* and +*2.4drl:EGFP (pDestTol2pA2_ +2.4drl:EGFP)* were assembled from pENTR/5’_+*2.0drl* or pENTR/5’_+*2.4drl* together with *pME-β-globin_minpromoter_EGFP* (Tamplin et al., 2015), Tol2kit vectors *#302* (*p3E_SV40polyA*), and *#394* (*pDestTol2A2*) as backbone (Kwan et al., 2007). +*2.0drl:creERT2 (pDestTol2CY_+2.0drl:creERT2,alpha-crystallin:YFP)* and - *5.0sox17:creERT2 (pDestTol2CY_-5.0sox17:creERT2,alpha-crystallin:YFP)* were assembled from *pENTR/5’_+2.0drl* or *pENTR/5’_-5.0sox17* together with *pCM293* (*pENTR/D_creERT2*) (Mosimann et al., 2011), Tol2kit vector *#302* (*p3E_SV40polyA*), and *pCM326 (pDestTol2CY)* as backbone (Mosimann et al., 2015). Genomic coordinates for the +*2.0drl* enhancer used in the described constructs are *chr5:61,649,227-61,650,194* (*GRCz11/danRer11*). Transcription factor binding sites were predicted using the JASPAR online interface (Khan et al., 2018).

The regulatory elements of mouse-specific LPM enhancers were PCR-amplified (Extended Data Table 1; regulatory elements), TOPO-cloned into the *pENTR5’* plasmid, and assembled together with *pKD001* (*pME-β-globin_minpromoter_EGFP* with improved Kozak sequence), Tol2kit vectors *#302* (*p3E_SV40polyA*), and *#394* (*pDestTol2A2*) as backbone.

Assembled reporter constructs were injected at a concentration of 25 ng/μl together with 25 ng/μl *Tol2* mRNA for Tol2-mediated zebrafish transgenesis (Felker and Mosimann, 2016). Injected F0 founders were screened for specific EGFP or *alpha-crystallin:*YFP expression. Zebrafish were raised to adulthood and screened in F1 for germline transmission. Single-insertion transgenic strains were generated, and microscopy images were taken on a Leica M205FA with a Leica DFC450C digital camera or a Leica SP8 confocal microscope with Plan-Apochromat 20x/0.5 objective. Images were processed using Leica LAS and Fiji (Schindelin et al., 2012).

Established transgenic and mutant lines used in this study included *drl:EGFP* (Mosimann et al., 2015), *drl:mCherry^zh705^* (Sánchez-Iranzo et al., 2018), *drl:creERT2* (Mosimann et al., 2015), *ubi:creERT2* (Mosimann et al., 2011), *ubi:switch* (Mosimann et al., 2011), *hsp70l:Switch^zh701^*(Felker et al., 2018), *lmo2:dsRED2* (Zhu et al., 2005), *scl/tal1:EGFP* (Jin et al., 2006), *pax2.1:EGFP* (Picker et al., 2002), *hand2:EGFP* (Yin et al., 2010), *actb2:h2afva-mCherry* (Krens et al., 2011), maternal and zygotic EGF-CFC co-receptor *oep* gene mutants (*MZoep*) (Gritsman et al., 1999), and maternal-zygotic *somitabun^dtc24^* mutants (*sbn*, dominant-negative *smad5*) (Hild et al., 1999; Mullins et al., 1996).

### Zebrafish CreERT2-based lineage tracing

Lineage tracing experiments were performed by crossing female *hsp70l:Switch* or *ubi:Switch* (Mosimann et al., 2011) reporter carriers with the male *creERT2* drivers *drl:creERT2* (Mosimann et al., 2015), *sox17:creERT2*, and +*2.0drl:creERT2*. Embryos were induced using 4-Hydroxytamoxifen (4-OHT) (H7904; Sigma H7904) from fresh or pre-heated (65°C for 10 min) stock solutions in DMSO at a final concentration of 10 μM in E3 embryo medium (Felker et al., 2016) at indicated time points. Embryos were washed in untreated E3 medium at 24 hpf. To induce EGFP transcription in *hsp70l:Switch* carrying embryos, the embryos were incubated at 37°C for 1 hour, 3 hours before fixation. Embryos were fixed with 4% paraformaldehyde (PFA) overnight at 4°C at 3 dpf and processed for confocal analysis.

### Zebrafish transverse vibratome sections

Transverse sections were generated as previously described (Gays et al., 2017). Fixed embryos were washed in PBS, embedded in 6% low-melting agarose (Sigma-Aldrich) in PBS/0.1% Tween-20 (Sigma-Aldrich), and cut into 130 μm thick sections using a vibratome (Leica VT 1000S). Sections were mounted in DAPI-containing Vectashield (Cat#H-1200; Vector Laboratories). Sections were analyzed with a Zeiss LSM710 confocal microscope with a Plan-Apochromat 40x/1.3 oil DIC M27 objective. Images were cropped and adjusted for brightness using ImageJ/Fiji (Schindelin et al., 2012). Graphs were generated in Graphpad Prism 5.

### Zebrafish morpholino and crispant experiments

The previously established *sox32* morpholino (Sakaguchi et al., 2001) was synthesized by GeneTools (sequence: *5’-TGCTTTGTCATGCTGATGTAGTGGG-3’*) and dissolved in nuclease-free water to a stock concentration of 10 µg/µl. The *sox32* morpholino injection mix of 4 µg/µl was incubated for 10 min at 65°C, and 8 ng were microinjected into one-cell stage zebrafish embryos.

sgRNAs were obtained in an oligo-based approach as previously described (Burger et al., 2016). Briefly, primer extension was performed using Phusion polymerase (NEB) of forward primer 5’-*GAAATTAATACGACTCACTATAGG-N_20_-GTTTTAGAGCTAGAAATAGC*-3’ (including a 20 nt target site) and invariant reverse primer 5’-*AAAAGCACCGACTCGGTGCCACTTTTTCAAGTTGATAACGGACTAGCCTTATTTTAACTTGCTA TTTCTAGCTCTAAAAC*-3’ (PAGE-purified) (Bassett et al., 2013). The following target sites are shown in this manuscript: 5’-*AGATTTGTTTAGTCAGTGTC*-3’ (*drl ccG*), 5’-*GTCTGGAAACAGTCTGAATC*-3’ (*drl ccH*), 5’-*GAGACTTCGCCCTTCGGTTC*-3’ (*mixl1 ccB*), and 5’-*GACAGAACAGGCCACGTTGA*-3’ (*mezzo ccA*).

*In vitro* transcription of sgRNAs was performed as previously described (Burger et al., 2016) using MAXIscript T7 (Ambion). Afterwards, RNA was precipitated with ammonium acetate, washed with 75% ethanol, and dissolved in DEPC water. RNA quality was checked on a MOPS gel. RNPs of Cas9 protein (Cas9-mCherry, *pMJ293* (Burger et al., 2016) (available from Addgene)) and sgRNA were *in vitro*-assembled for 10 min at 37°C in 300 mM KCl to ensure maximum cleavage efficiency and microinjected into the cell of one-cell stage embryos (Burger et al., 2016). The CRISPR target regions in the transgenic locus were amplified using specific target region amplification primers (Table S1; target region amplifications).

Sequencing analysis was performed with the R software package CrispRVariants for allele- and sequence-level variant counting and visualization (Lindsay et al., 2016).

### Zebrafish overexpression experiments

For early developmental transcription factor genes, ORFs were PCR-amplified from mixed-stage zebrafish cDNA using ORF-specific primers (Table S1; coding sequence primers); full-length zebrafish *eomesA* was derived from I.M.A.G.E clone *IRBOp991A0739D* (Source BioScience LifeSciences, UK). All CDS were TOPO-cloned into the *pENTR/D™* Directional TOPO® plasmid (Invitrogen) according to the manufacturer’s instructions. Constitutive-active Smad2, which encodes an N-terminal truncation of zebrafish Smad2 (Dick et al., 2000), was amplified from zebrafish cDNA and subcloned into *pENTR1A* (*pCM269*). Subsequent cloning reactions were performed with the Multisite Gateway system with LR Clonase II Plus (Invitrogen) according to the manufacturer’s instructions and as previously described (Felker and Mosimann, 2016). The *CDS* as pENTR/D vectors (*pENTR1A* for *smad2(ca)*) were assembled with *pENTR5’_ubi* (Mosimann et al., 2011), *pENTR5’_T7-VP16*, *pENTR5’_T7-eng* or *pENTR5’_T7* together with Tol2kit vectors *#302* (*p3E_SV40polyA*), and *#394* (*pDestTol2A2*) as backbone (Kwan et al., 2007). Afterwards, plasmids were linearized (in case of *T7-VP16, T7-eng*, or *T7*) and *in vitro* transcribed (Roche). The *EomesA-VP16* plasmid (containing 153aa-431aa of the zebrafish EomesA ORF) (Bruce et al., 2003) was kindly provided by Dr. Rebecca Burdine. The plasmid was linearized via NotI followed by SP6 *in vitro* transcription. The *Ntl-VP16* plasmid was kindly provided by Dr. Ashley Bruce. The *etv5a-VP16* and *etv5a-eng* plasmids were kindly provided by Dr. Michael Tsang. All mRNAs were precipitated with ammonium acetate, washed with 75% ethanol, and dissolved in DEPC water. mRNA quality was checked on MOPS gels as described (Burger et al., 2016).

### Zebrafish chemical treatments

Chemicals for performed zebrafish treatments were dissolved in DMSO. Dorsomorphin (10-30 µM; Sigma-Aldrich) (Yu et al., 2008) and SB-505124 (30-60 µM; Sigma-Aldrich) (DaCosta Byfield et al., 2004; Rogers et al., 2017) were administered at 1-cell stage and embryos kept in the treated E3 until fixation.

### Zebrafish whole-mount in situ hybridization

Total RNA was extracted from zebrafish embryos from various stages during development. This RNA was used as template for generation of first-strand complementary DNA (cDNA) by the Superscript III First-Strand Synthesis kit (Cat#18080051; Invitrogen). *In situ* hybridization (ISH) probes were designed with an oligonucleotide-based method (including T7 promoter added to the reverse primers) using zebrafish cDNA (Table S1; *in situ* hybridization probes). The following oligonucleotide pairs (including T7 promoter added to the reverse primers) were used to amplify the DNA template from zebrafish cDNA. The ISH probe for *admp* was obtained from a pCS2_ADMP plasmid, and *sizzled* from *pCS2_Sizzled*. *Admp* and *sizzled* were linearized by ClaI. The *gata2a* probe was obtained from the middle entry vector *pCM238*. For *in vitro* transcription, T7 RNA polymerase (Roche) and digoxigenin (DIG)-labeled NTPs (Roche) were used. Afterwards, RNA was precipitated with lithium chloride, washed with 75% ethanol, and dissolved in DEPC water. RNA quality was checked on a MOPS gel. ISH on whole-mount zebrafish embryos was executed as described before (Thisse and Thisse, 2008). After ISH, embryos were transferred to 80-95% glycerol (Sigma-Aldrich), and microscopy images were taken on a Leica M205FA with a Leica DFC450C digital camera. Images were cropped and adjusted for brightness using ImageJ/Fiji. The pathway schematic in Fig 4A was generated using BioRender.

### Zebrafish selective plane illumination microscopy (SPIM)

At 30-50% epiboly, embryos in the chorion were embedded into 1% low-melting agarose with optional 0.016% Ethyl 3-aminobenzoate methanesulfonate salt (Tricaine, Cat#A5040; Sigma) in E3 embryo medium, and sucked into an FEP tube (inner diameter: 2.0 mm, wall thickness: 0.5 mm). 6-7 embryos were positioned on top of each other. The FEP tube was mounted in the microscope imaging chamber filled with E3 medium. Panoramic (3D) SPIM/lightsheet microscopy and subsequent image processing (Mercator projections) were performed as previously described (Schmid et al., 2013). A z-stack of 402 planes was obtained from every embryo with an interval of 2 min for a period of 14-17 hours. Images were processed using Leica LAS, ImageJ/Fiji, and Photoshop CS6.

### Chicken embryo incubation and ex-ovo culturing

*Ex ovo* culturing was adapted from previously established protocols (New, 1955). Fertilized chicken eggs were obtained from a local hatchery and stored at 12°C up to maximum of 14 days. Prior to use, eggs were incubated horizontally for 17 hours until Hamburger-Hamilton (HH) 3+/4 in a 39°C incubator with 55-65% humidity. After incubation, the eggs were kept for at least 30 min at RT before opening. Eggs were opened in a petri dish and a layer of thick albumin together with the chalaziferous layer was removed using a plastic Pasteur pipette. A paper ring was placed around the embryo on the yolk and dissection scissors were used to cut the yolk membrane around the ring. The paper ring with the embryo was cleaned from remaining yolk and transferred and placed upside down on a semisolid albumin/agarose (43.5 ml thin albumin incubated for 2 hours at 55°C, 5 ml 2% agarose, 1.5 ml 10% glucose in 30 mm petri dishes) culturing plate. Embryos were recovered for at least 2 hours at RT before electroporation.

### Chicken embryo injection and electroporation

For electroporations, a customized electroporation chamber was used containing an electrode with a positive pole on the bottom of the chamber and separate negative electrode on a holder (kindly provided by the lab of Dr. Jerome Gros). Both electrodes were mounted and connected to a square wave electroporator (BTX ECM 830). The electroporation chamber was filled with HBSS (Gibco Life Technologies), and the embryo-containing paper ring was placed in the chamber with the dorsal side up. The DNA mixture was injected by a mouth injector along the primitive streak beneath the pellucid membrane. The positive electrode holder was placed on top of the streak to allow electricity pulses flow through the embryo (3 pulses, 8V x 50 milliseconds, 500 milliseconds interval). The embryos were placed back on the albumin culturing plates with the ventral side up and placed back at 39°C until HH8-9. Microscopy images of the embryos were taken at HH8-9 on a Leica M205FA with a Leica DFC450C digital camera. Images were cropped and adjusted for brightness using ImageJ/Fiji. All injection mixtures for electroporations contained 0.1% fast green dye, 0.1% methyl-cellulose, 300 ng/µl control plasmid *pCAGGs* (*pCMV:H2B-CAGG-RFP*, abbreviated for *chicken β-actin promoter CAGG-mCherry)* and 1 µg/µl of the plasmid of interest.

### Axolotl experiments

The generation of transgenic animals and determination of developmental stages were performed as described previously (Bordzilovsakya et al., 1989; Khattak et al., 2014). Animals at stage 43 were anaesthetized by bathing in 0.01% benzocaine (Khattak et al., 2014).

Live imaging was performed on an Olympus SZX16 fluorescence stereomicroscope. Time lapse movies were acquired using an Axio Zoom.V16 (Zeiss) stereomicroscope. Confocal images were acquired on a Zeiss LSM780-FCS inverted microscope.

For immunostaining, embryos were fixed in MEMFA at 4°C overnight, washed in PBS, embedded in 2% low melting temperature agarose and sectioned by vibratome into 200 µm thick sections. Fibronectin was detected using mouse anti-Fibronectin antibody (ab6328, Abcam) at 5µg/ml.

### Lamprey experiments

The +*2.0drl* regulatory element was amplified from the zebrafish vector +*2.0drl:EGFP* by PCR using KOD Hot Start Master Mix (Novagen) (Table S1; regulatory elements). The amplified enhancers were cloned into the HLC vector for lamprey transgenesis (Parker et al., 2014b), containing the mouse *c-Fos* minimal promoter, by Gibson assembly using the Gibson Assembly Master Mix (NEB). Injections for I-SceI meganuclease-mediated lamprey transient transgenesis were performed using *P. marinus* embryos at the one-cell stage with injection mixtures containing 0.5 U/µl I-SceI enzyme and 20 ng/µl reporter construct as described previously (Parker et al., 2014a). Selected EGFP-expressing embryos were fixed in MEM-FA and dehydrated in methanol for *in situ* hybridization. EGFP-expressing embryos were imaged using a Zeiss SteREO Discovery V12 microscope with variable zoom and a Zeiss Axiocam MRm camera with AxioVision Rel 4.6 software. Images were cropped and adjusted for brightness using Adobe Photoshop CS5.1.

For mRNA ISH, total RNA was extracted from st 21-26 *P. marinus* embryos using the RNAqueous Total RNA Isolation Kit (Ambion). This was used as a template for 3’ rapid amplification of cDNA ends (RACE) with the GeneRacer Kit and SuperScript III RT (Invitrogen). A 339bp-long *pmHandA in situ* probe was designed based on a characterized cDNA sequence from the closely related Arctic lamprey (*Lethenteron camtschaticum*) (Kuraku et al., 2010), and this sequence was amplified by PCR from 3’ RACE cDNA using KOD Hot Start Master Mix (Novagen) with the following primers: 5’-*GCGGAGGACATTGAGCATC*-3’ (forward) and 5’-*TGGAATTCGAGTGCCCACA*-3’ (reverse). The cDNA fragment was cloned into the *pCR4-TOPO* vector (Invitrogen). The 709bp-long *eGFP* probe was described previously (Parker et al., 2014b).

DIG-labelled probes were generated and used in lamprey whole-mount ISH as described previously (Sauka-Spengler et al., 2007). Embryos were cleared in 75% glycerol and imaged using a Leica MZ APO microscope with variable zoom and Lumenera Infinity 3 camera with Lumenera Infinity Capture v6.5.3 software. Images were cropped and adjusted for brightness using Adobe Photoshop CS5.1.

After ISH, selected embryos were transferred into 30% sucrose in PBS, embedded in O.C.T. Compound (Tissue-Tek), and cut into 10 µm-thick cryosections using a CryoStar NX70 cryostat (Thermo Scientific). Images were taken using a Zeiss Axiovert 200 microscope with an AxioCam HRc camera and AxioVision Rel 4.8.2 software. Original data underlying the lamprey experiments in this manuscript are accessible from the Stowers Original Data Repository at http://odr.stowers.org/websimr/.

### Ciona experiments

+*2.0drl* was amplified from the zebrafish vector +*2.0drl:EGFP* and sub-cloned upstream of *unc76:GFP* to generate a Ciona reporter construct including minimal promoter (pBuS24; see Table S1, regulatory elements for primer sequences). Gravid *Ciona robusta* adults were obtained from M-REP (San Diego CA, USA). To test the activity of the zebrafish enhancers in *Ciona robusta*, 80 μg of +*2.0drl:EGFP* was injected in a mixture with the reporter plasmid for *Mesp* (Davidson et al., 2005) to mark the B7.5 cardiopharyngeal lineage with *H2B:mCherry* (10 μg). For antibody staining, embryos were fixed in 4% MEM-PFA for 30 min, rinsed several times in PBT (PBS/0.1% Tween-20), and incubated with anti-GFP (1:500, mouse mAb, Roche) with 2% normal goat serum in PBT at 4°C overnight. Embryos were washed in PBT and then incubated with donkey anti-mouse secondary antibody (1:1000) coupled to Alexa Fluor 488 (Life Technologies) in PBT with 2% normal goat serum for 2 hours at RT, then washed in PBT (Racioppi et al., 2014).

### Amphioxus experiments

The regulatory elements *drl* (entire 6.35 kb) and +*2.0drl* were amplified from the zebrafish reporter vector *drl:EGFP* (Mosimann et al., 2015) and +*2.0drl:EGFP* and subcloned upstream of a EGFP reporter in the *pPB* vector carrying *PiggyBac* transposon terminal repeats (Kozmikova and Kozmik, 2015). Adults of *Branchiostoma lanceolatum* were collected in Banyuls-sur-mer, France, prior to the summer breeding season and raised in the laboratory until spawning. The spawning of amphioxus male and females was induced by shifting of the temperature as described (Fuentes et al., 2007). For microinjection of amphioxus eggs, a mixture of *pPB-drl:EGFP* or *pPB-+2.0drl:EGFP* (200 ng/μl) with PiggyBac transposase mRNA (100 ng/μl) in 15% glycerol was used. Transgenic embryos were allowed to develop until neurula stage, fixed in 4% PFA overnight at 4°C, mounted with Vectashield with DAPI (Vector Laboratories), and analyzed using a Leica SP5 confocal microscope. The confocal images were adjusted for brightness and contrast with ImageJ/Fiji.

---

**Table S1:**
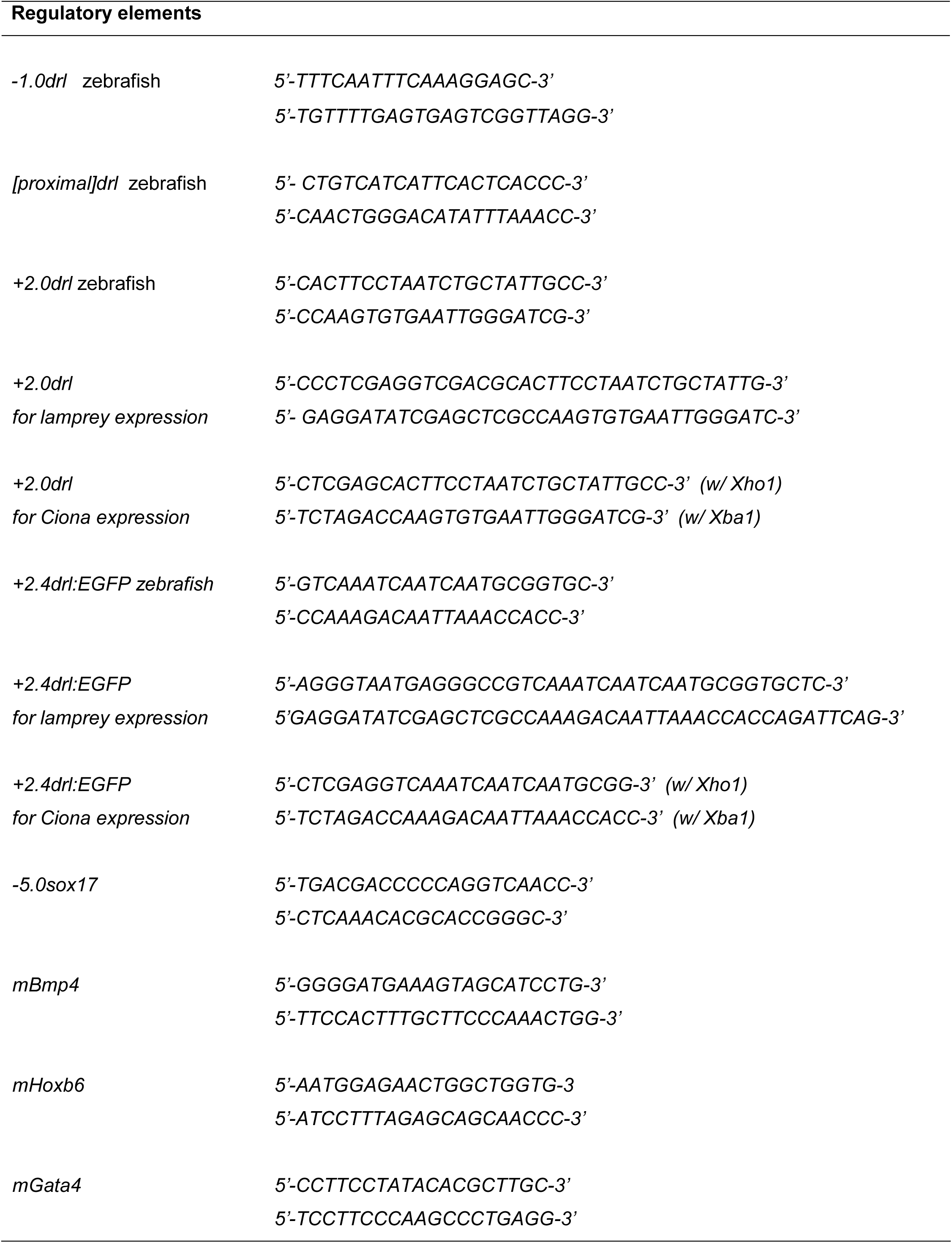
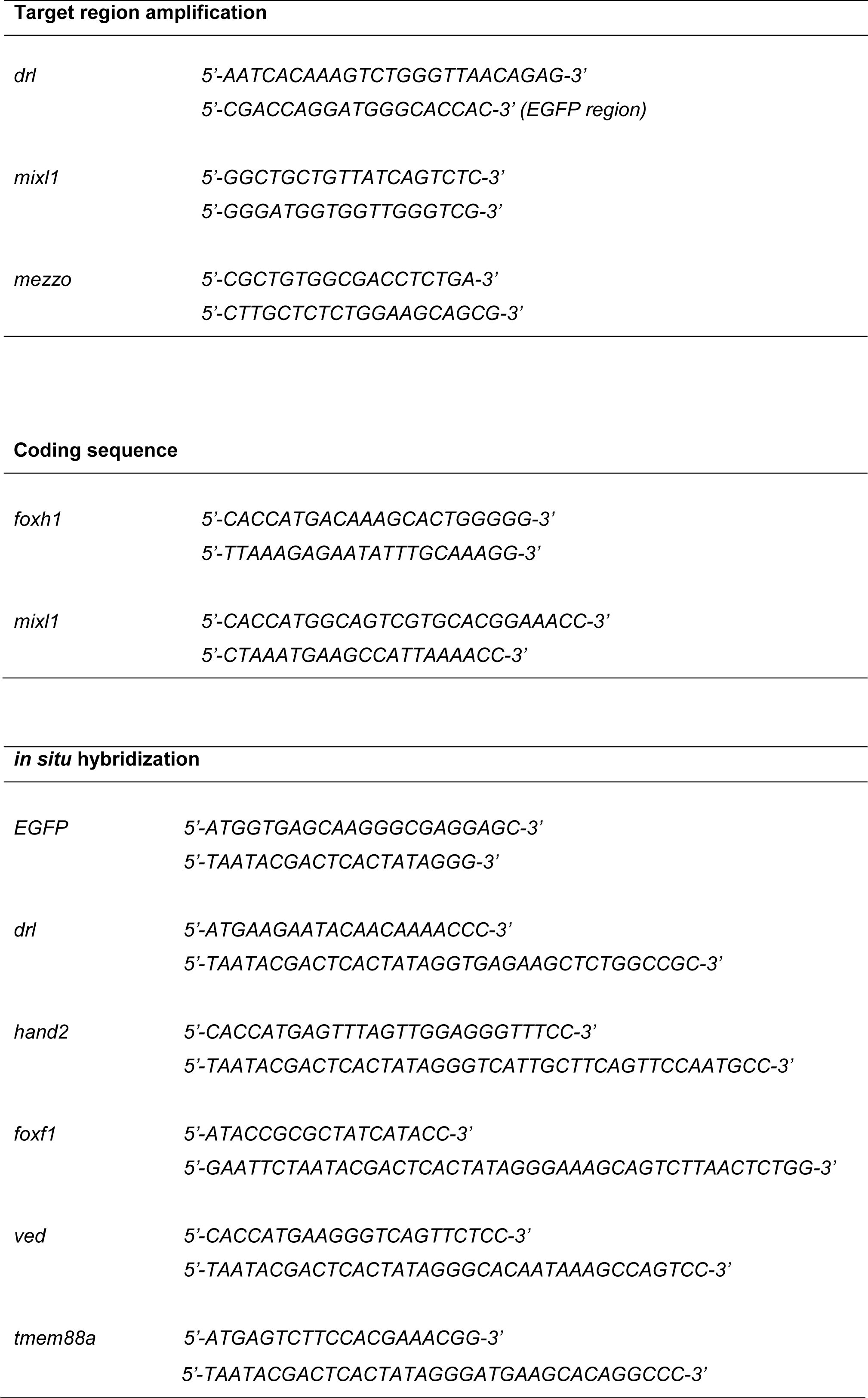
Primer sequences.

**Fig. S1:**
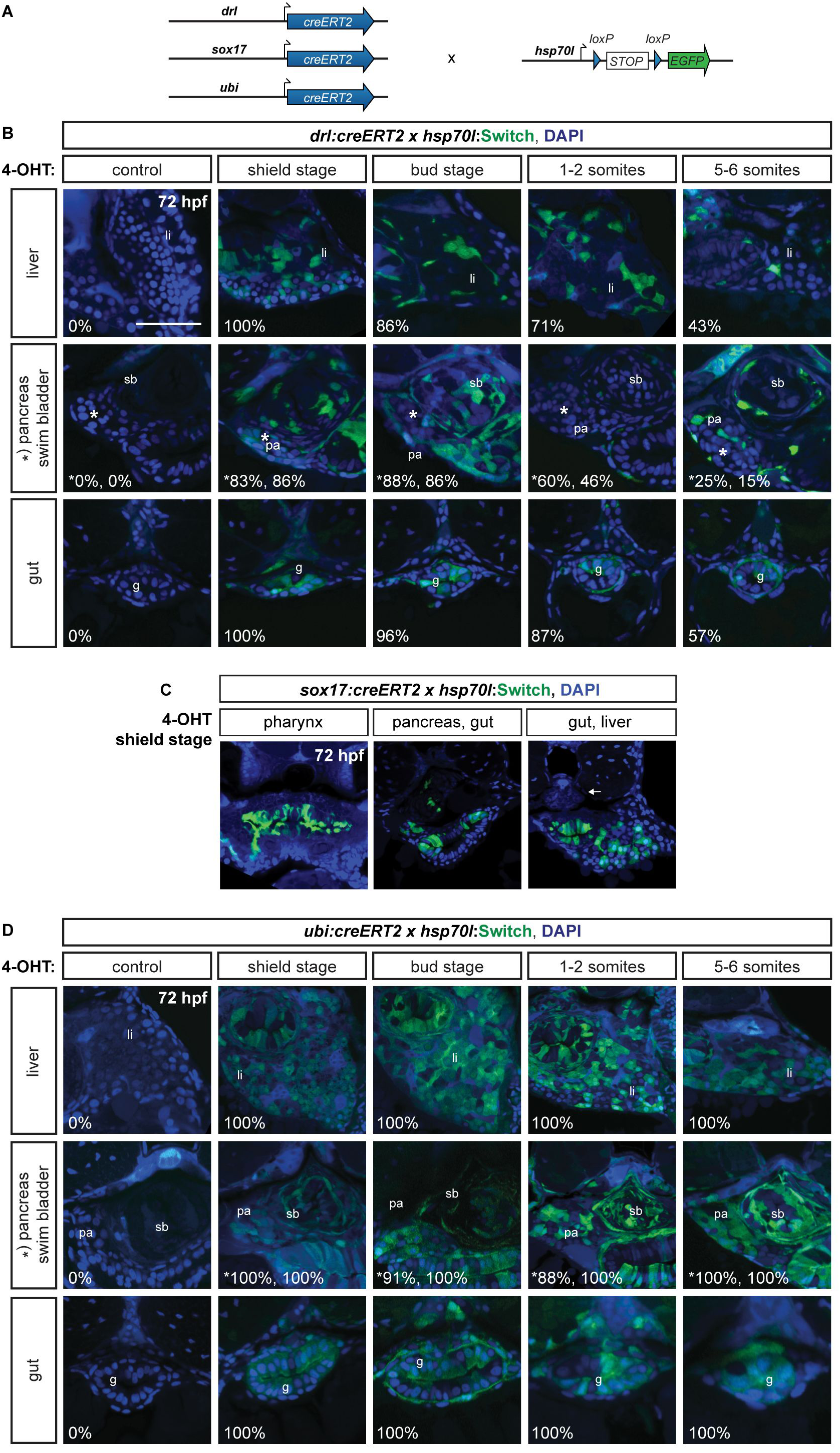
*drl* reporter-expressing cells contribute to mesendoderm with LPM-restricted mesoderm contribution. (**A**) Schematic representation of *drl:creERT2*, *sox17:creERT2*, and *ubi:creERT2* crosses to *hsp70l:Switch* for genetic lineage tracing. (**B-D**) Whole-body transverse sections of lineage traced embryos at 72 hpf following different 4-OHT induction time points (shield, tailbud, 1-2 ss, and 5-6 ss) to trigger *hsp70l:Switch* recombination (green), nuclei counterstained with DAPI (blue). Numbers indicate the percentage of embryos showing lineage-labeling in depicted organs. (**B**) *drl:creERT2* with 4-OHT induction at shield stage traces, besides of LPM-derived tissue, a high percentage of endoderm-derived lineages, as depicted for liver (li), swim bladder (sb), pancreas (pa), and gut (g), while 4-OHT induction at 5-6 ss traces shows reduced to minimal lineage labeling in endoderm-derived tissue. (**C**) Transverse sections of *sox17:creERT2* lineage-traced endoderm cells after 4-OHT induction at shield stage. Lineage labeling is confined to endodermal lineages and absent from mesodermal and specifically LPM lineages, as depicted for the pharynx, pancreas, gut, and liver, in contrast to LPM-derived tissue such as the aorta (arrow). (**D**) No tissue bias or obvious absence of EGFP in lineage reporter labeling could be detected for the *hsp70l:Switch* line, as shown using the ubiquitous *ubi:creERT2*. Scale bar 50 μm.

**Fig. S2:**
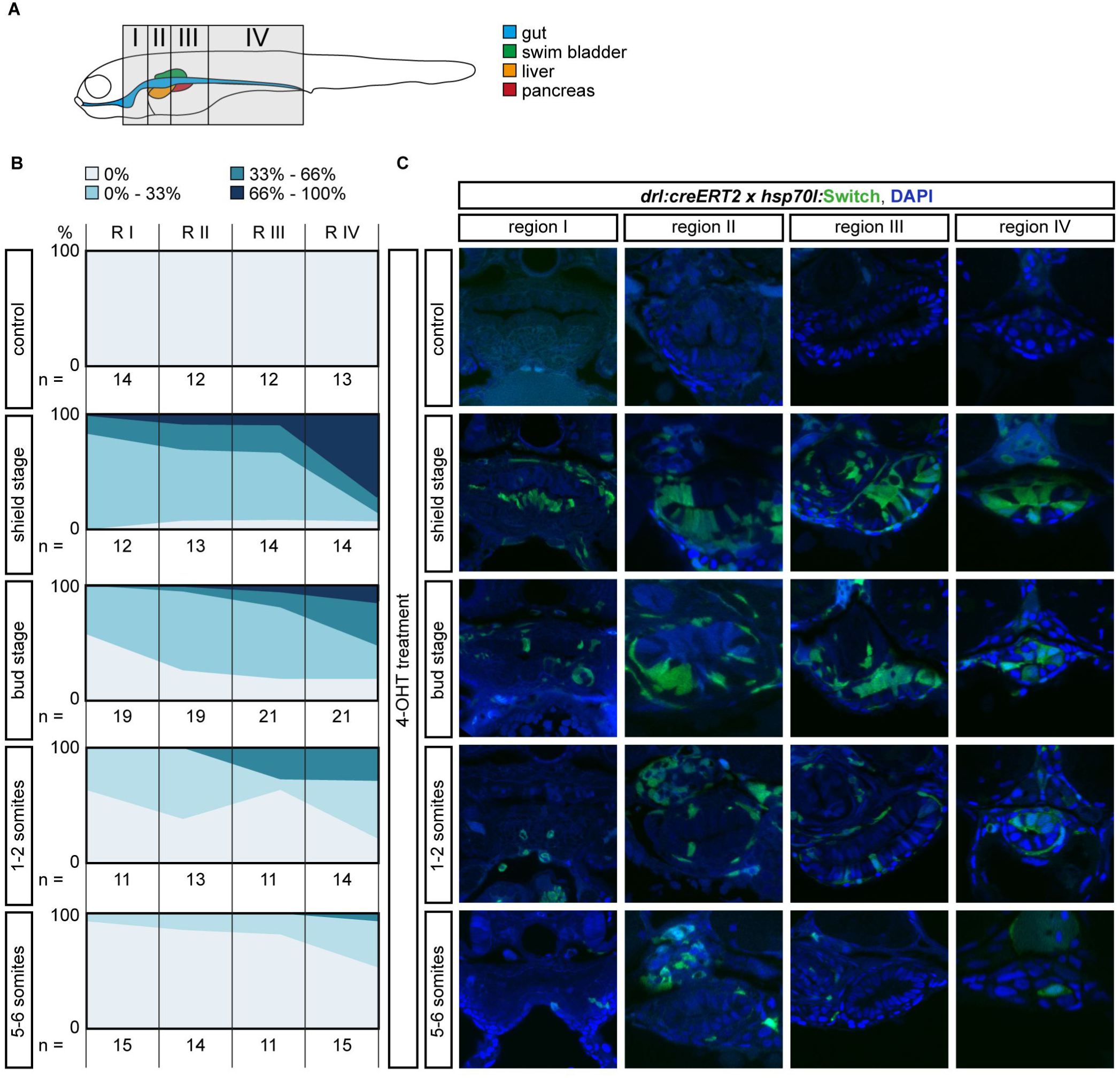
The LPM specifies within the mesendoderm with an anterior-posterior gradient in - *6.35drl* lineage traced embryos. **(A)** Schematic of the four regions I-IV defined to study anterior-posterior differences in lineage-labeling of the gut epithelium. Region I is the most rostral and includes the pharynx and the heart; Region II inlcudes the esophagus, the beginning of the swim bladder (pneumatic duct), the liver, and the anterior tip of the pancreas; Region III includes the gut, pancreas, and swim bladder; region IV includes the most caudal part of the gut and is recognizable by the yolk extension. **(B)** Transverse sections of the gut of 3 dpf *drl:creERT2;hsp70l:Switch* embryos. Columns represent the region and rows represent the stage of 4-OHT induction. **(C)** For each section, the switching efficiency in the gut epithelium was classified into 0%, between 0-33%, between 33-66% or between 66-100%. The Y-axis represents the fraction number of embryos. Indicated n-numbers represent the number of embryos analyzed. Embryos were collected in three independent experiments.

**Fig. S3:**
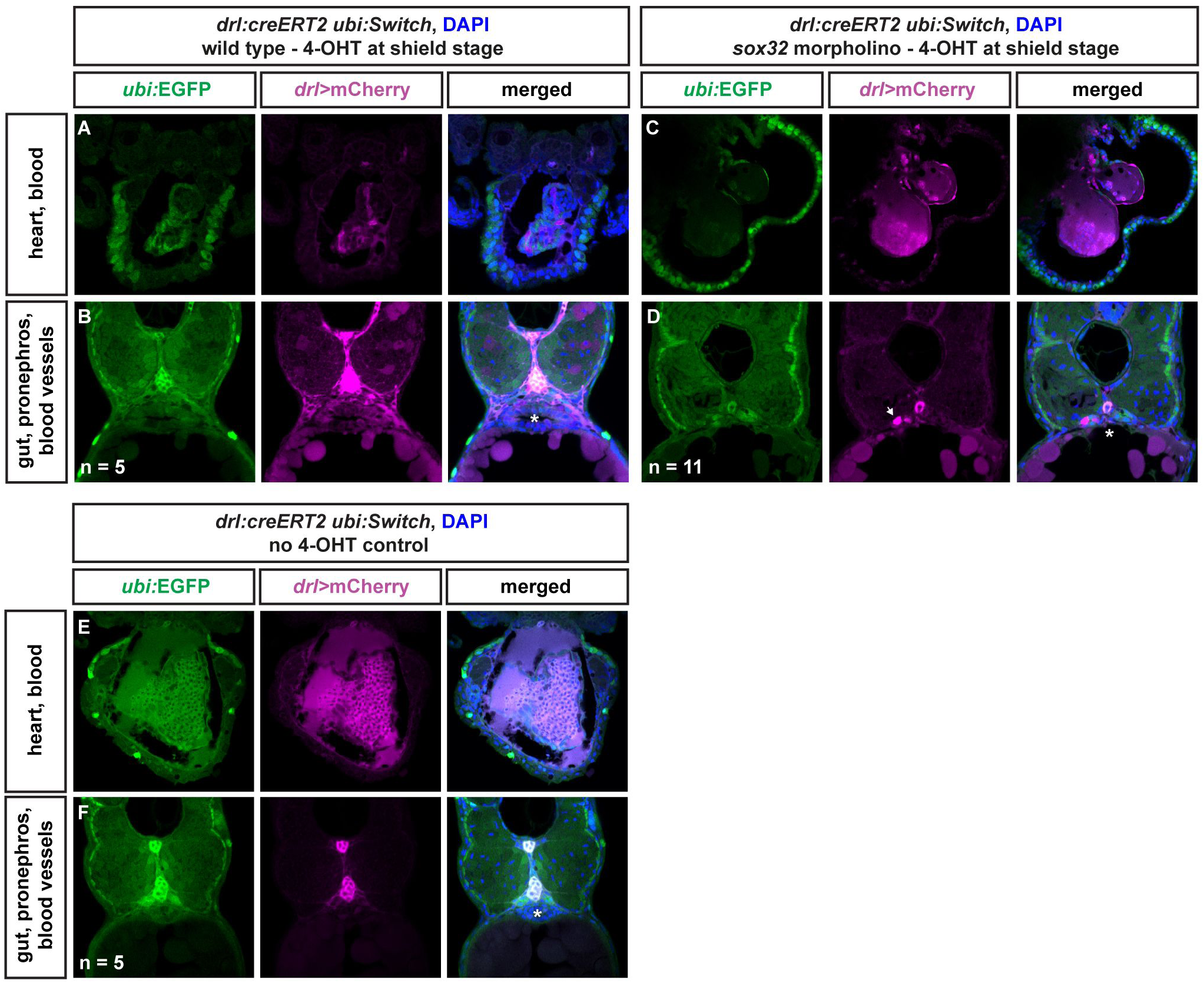
LPM lineages and organs still arise in embryos devoid of endoderm. Transverse sections of *drl:creERT2* x *ubi:Switch* (*ubi:lox-GFP-lox_mCherry*) embryos fixed at 3 dpf. (**A,B**) Embryos were induced with 4-OHT at shield stage, bringing *mCherry* under control of the *ubi* promoter in cells with active CreERT2 from shield stage (magenta), while unrecombined cells keep expressing GFP (green). LPM-derived organs, as depicted for heart, blood, pectoral fins, pronephros, and endothelial cells, and endoderm-derived lineages, including swim bladder, gut epithelium, and liver, are lineage-traced. (**C-D**) In *sox32* morphants that are devoid of endoderm, 4-OHT induction at shield stage still traced LPM-derived organs, as depicted for heart, blood, pectoral fins, pronephros (arrow), and endothelial cells. (**E-F**) Control without 4-OHT admission reveals absence of any background activity of the used *ubi:Switch* reporter (note autofluoresence of blood), confirming absence of leakiness of the *creERT* and *loxP* lines used. Asterisks indicate endoderm-derived gut epithelium (**B**, **F**) or absence thereof (**D**). Nuclei counterstained with DAPI (blue).

**Fig. S4:**
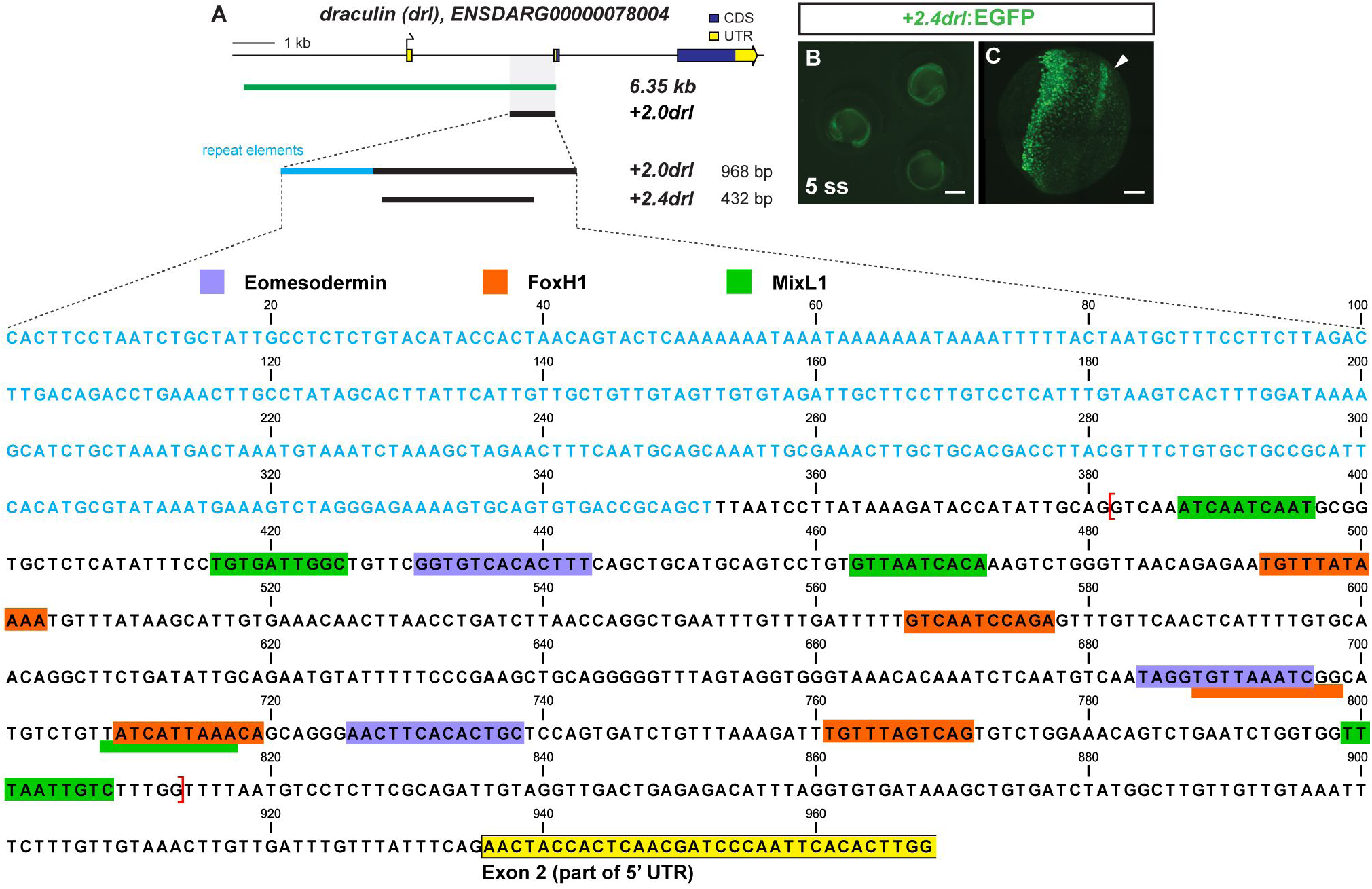
Trimming of +*2.0drl* to the minimal 432 bp LPM +*2.4drl* enhancer region. (**A**) Schematic representation of the −*6.35drl* locus including the PCR-based trimming approach to map the smallest LPM specific regulatory enhancer region; repetitive elements highlighted in blue. Sequence depicts the +*2.0drl* region in intron 1 (*Danio rerio* strain Tuebingen chromosome 5, GRCz11 primary assembly, *NC_007116.7:61649227-61650194*): blue text depicts repetitive elements, colored boxes depict predicted transcription factor binding sites (JASPAR database for vertebrates, 80-85% threshold), yellow frame marks the beginning of *drl* exon 2, and red brackets outline the sequence for the +*2.4drl* core sequence. (**B**) Expression of +*2.4drl:EGFP* at 5 ss; arrow indicates axial mesoderm expression, indicating increased promiscuity after trimming from the initial +*2.0drl* element. Scale bar 400 μm (**B**) and 80 μm (**C**).

**Fig. S5:**
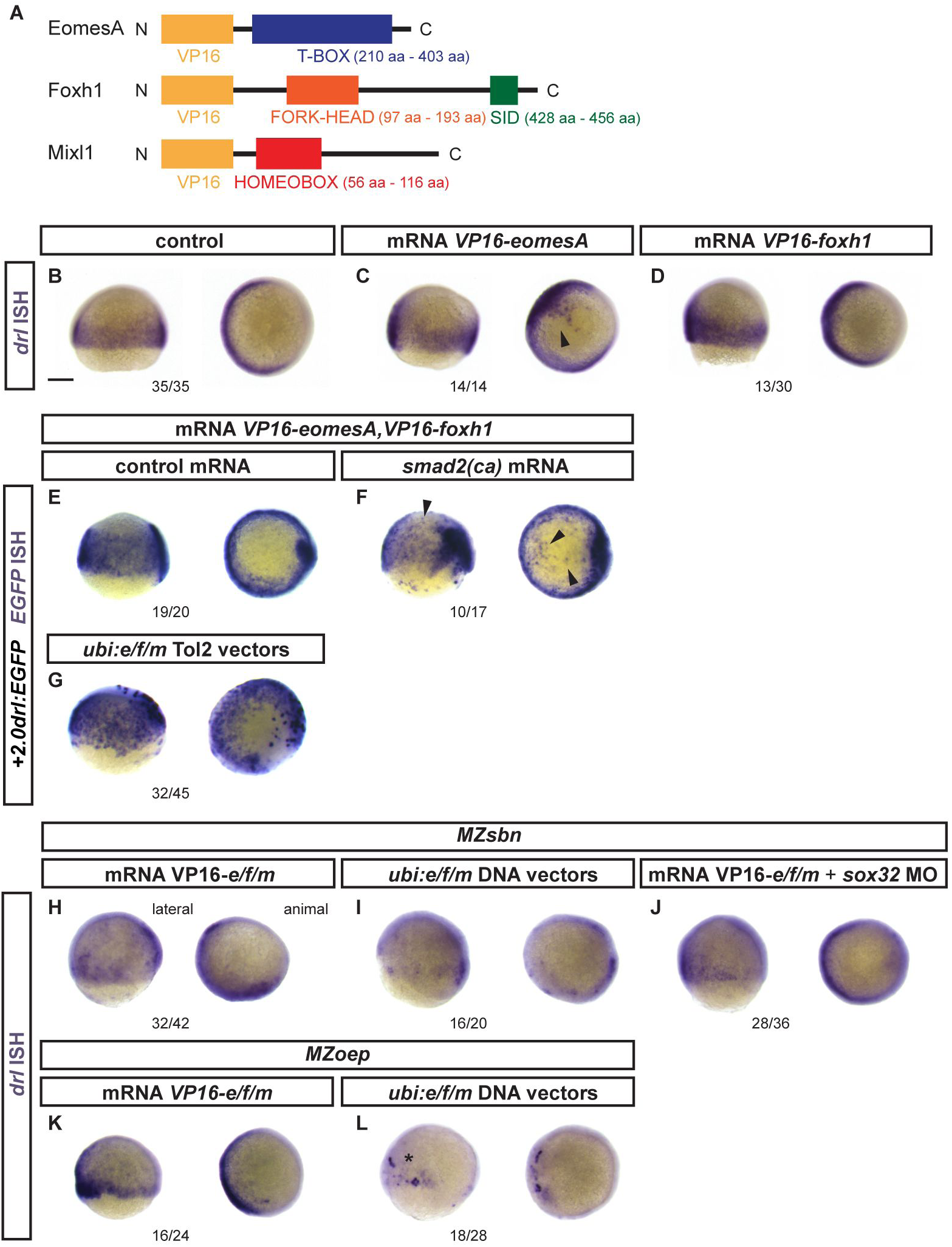
Response of endogenous *drl* and *drl* reporters to EomesA, Foxh1, and Mixl1 *expression*. (**A**) Schematics of EomesA, Foxh1, and Mixl1 fusion proteins with the N-terminally added VP16 transactivation domain to generate constitutively-active transcription factors. Note that the expressed VP16-EomesA is restricted to the T-box of EomesA. (**B-D**) mRNA ISH for endogenous *drl* expression in controls (**B**) and embryos injected with VP16-fusions based on *eomesA* (**C**) or *foxh1* (**D**). (**E-F**) mRNA ISH for *EGFP* transcript expression in +*2.0drl:EGFP* embryos at shield to 70% epiboly stage after injecting a combination of *VP16*-fused *eomesA* and *foxh1* (*e/f*) without (**E**) or with (**F**) constitutive-active *smad2* that mimicks pan-Smad signaling (*smad2(ca)*), resulting in dorsal widening of the reporter expression pattern. (**G**) Injection of Tol2-based constructs under ubiquitous *ubi* promoter control to drive native *eomesA* (full-length), *foxh1*, and *mixl1* (*e/f/m*), resulting in mosaic induction of +*2.0drl:EGFP*, including individual dorsal blastomeres. (**H-J**) Endogenous *drl* expression revealed by mRNA ISH in BMP signaling-mutant (*MZsbn*) embryos injected with either (**H**) *VP16-e/f/m* mRNA, (**I**) *ubi:e/f/m* DNA vectors, or (J) *VP16-e/f/m* mRNA in an endoderm-perturbed background (*sox32* morpholino). (**K,L**) mRNA ISH for endogenous *drl* expression in Nodal-mutant (*MZoep*) embryos injected with (**K**) *VP16-e/f/m* mRNA or (**L**) *ubi:e/f/m* Tol2 vectors; note the localized clones of strong *drl* upregulation on the ventral side upon *ubi:e/f/m* injection. Scale bar in (**B**) 250 μm.

**Fig. S6:**
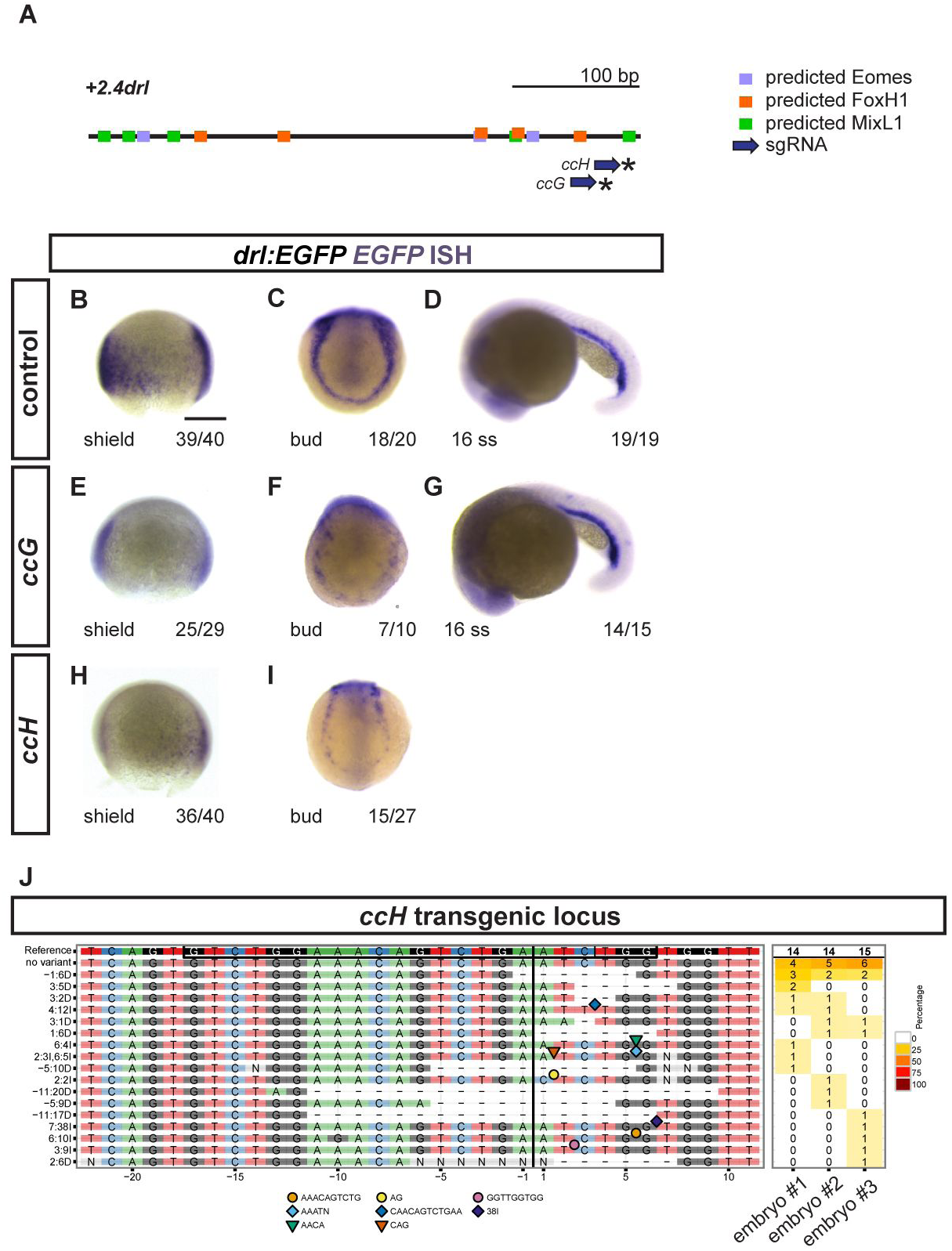
Crispant analysis of the +*2.0drl* region. (**A**) Schematic representation of the minimal +*2.4drl* LPM enhancer region (core of +*2.0drl*) with Eomes, FoxH1, and Mixl1 binding site predictions (JASPAR) and sgRNAs *ccG* and *ccH* annotated. (**B-I**) ISH for *EGFP* in +*2.0drl*:*EGFP* embryos shown for injection controls (**B-D**) or individually mutagenized using reconstituted Cas9 protein-sgRNA complexes with either sgRNA (**E-G**) *ccG* or (**H,I**) *ccH* (asterisks in **A**), both showing mosaic loss or reduction of reporter expression at shield stage compared to uninjected controls. Reporter expression that is dependent on different regulatory elements remains intact upon mutagenesis (**D,G**), indicating no overt mutagenesis of essential parts in the transgene itself. (**J**) Panel plot showing mutagenesis efficiency and allele spectrum in three representative *ccH* crispants, as created by CrispRVariants. The genomic reference sequence is shown on top, with the 20 bp *ccH* sgRNA followed by a 3 bp PAM indicated by the boxed regions. The Cas9 cleavage site is represented by a black vertical bar. Deletions are indicated by a ‘-‘, and insertion sequences by symbols. The right column of the plot shows detected allele frequency per analyzed embryo. Scale bar (**B**) 250 μm.

**Fig. S7:**
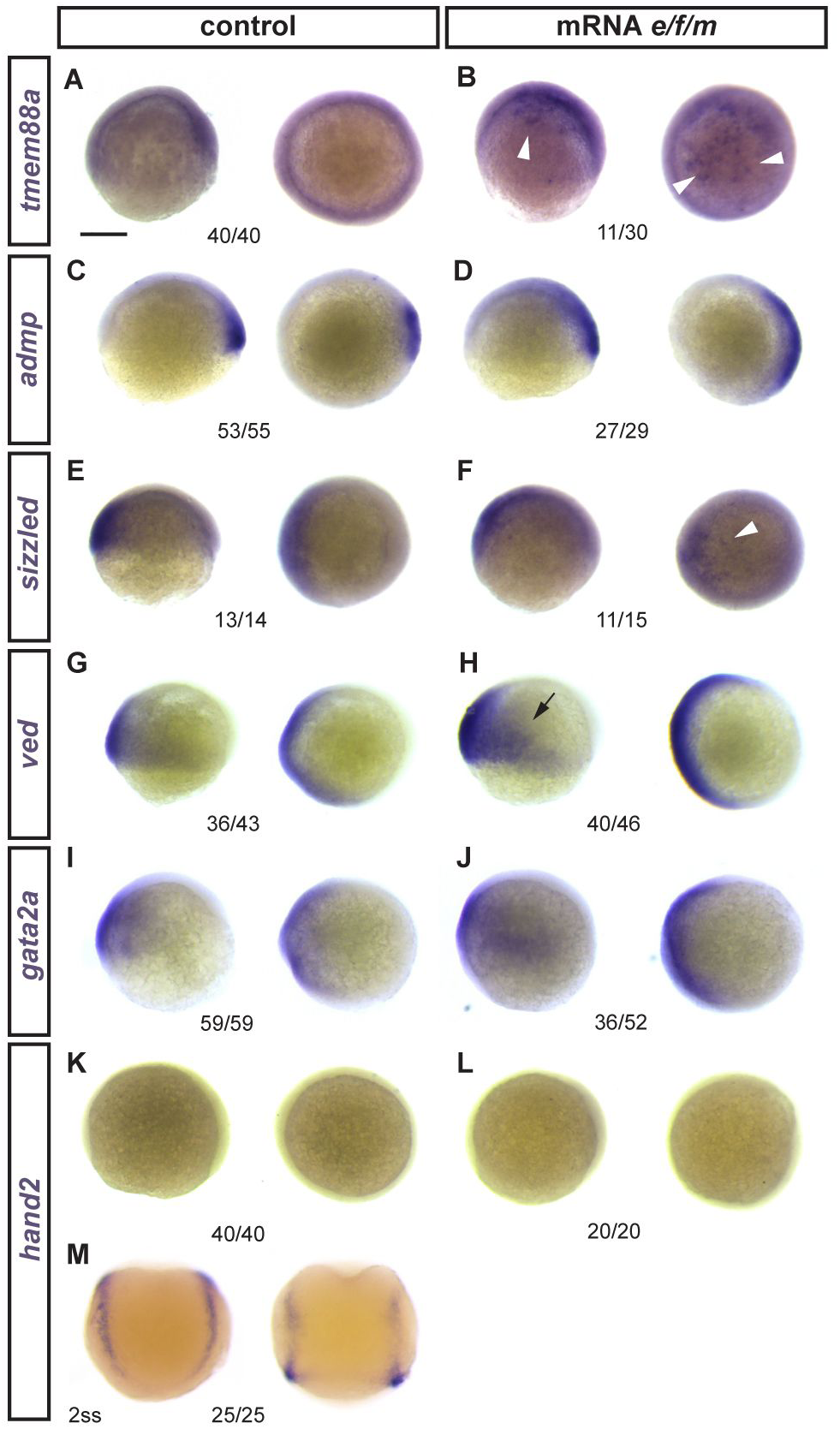
Effects of *eomesA, foxh1, and mixl1* misexpression on early developmental genes. (**A-M**) Zebrafish embryos ((**A-L)** shield to 75% epiboly or (**M**) 2 somite stage) as control or injected with mRNA for *eomesA*, *foxh1*, and *mixl1* (*e/f/m*), with mRNA ISH for indicated candidate genes. Latera view left, animal view right for each condition. (**A,B**) The early LPM gene *tmem88a* that is weakly expressed during epiboly (**A**) becomes activated in patches upon e/f/m misexpression (**B**, arrowheads). (**C-J**) Expression of various gastrulation-stage marker genes: *admp* (**C**,**D**), *sizzled* (slightly increased expression ventral, indicated by arrowhead) (**E**,**F**), *ved* (slight increased expression ventral, arrow) (**G**,**H**), *gata2a* (slight ventral expansion) (**I**,**J**). (**K-M**) The somite-stage LPM gene *hand2* does not become ectopically induced upon *e/f/m* misexpression (**K-L**), as tested with a functional mRNA ISH probe (shown for 2 somite stage in **M**). Scale bar (**A**) 250 μm.

**Fig. S8:**
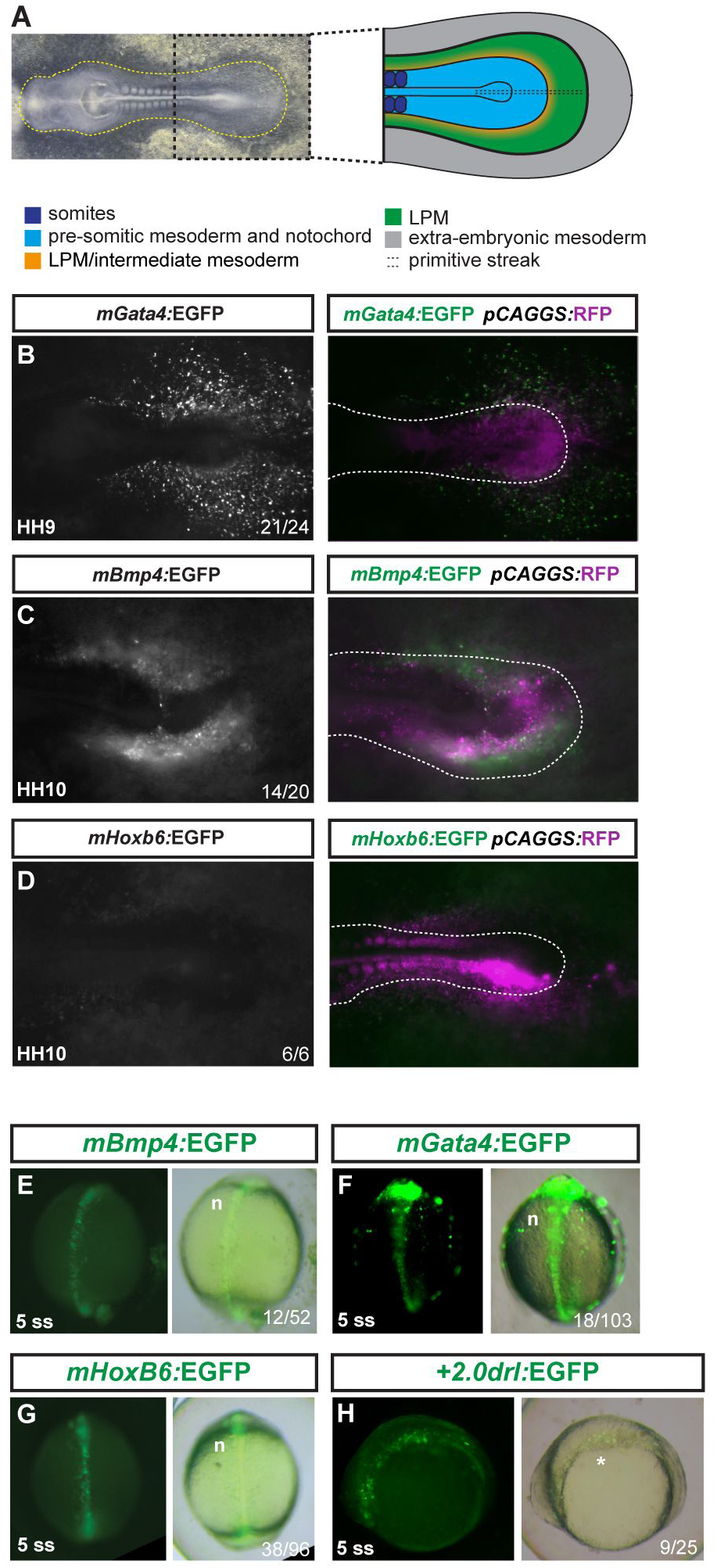
Mouse LPM enhancers show specific activity in chick, but not in zebrafish embryos. (**A**) Bright field image of an HH10 stage chicken embryo, anterior to the left, with schematic depiction of the posterior body (right), prospective LPM in green. (**B-D**) *Ex ovo*-cultured chicken embryos at HH9-HH10, electroporated at HH3+/H4 with reporters based on mouse LPM enhancers from *mGata4* (**B**), *mBmp4* (**C**), and *mHoxb6* (**D**) driving *EGFP* (grayscale), including a merged overlay together with electroporation marker plasmid *pCAGGS:RFP* (magenta, driving ubiquitous expression as electroporation control). Dashed lines (**B-D**) indicate the posterior outline of the individual chicken embryos, marking the predicted boundary between embryo and extra-embryonic tissue. (**E-H**) Zebrafish embryos depicted at early somitogenesis stages (dorsal views in **E-G**, lateral view in **H**) injected with EGFP reporters for *mBmp4* (**E**), *mGata4* (**F**), and *mHoxb6* (**G**) compared to +*2.0drl:EGFP* (**H**), revealing axial mesoderm expression of the tested mouse enhancers.

**Fig. S9:**
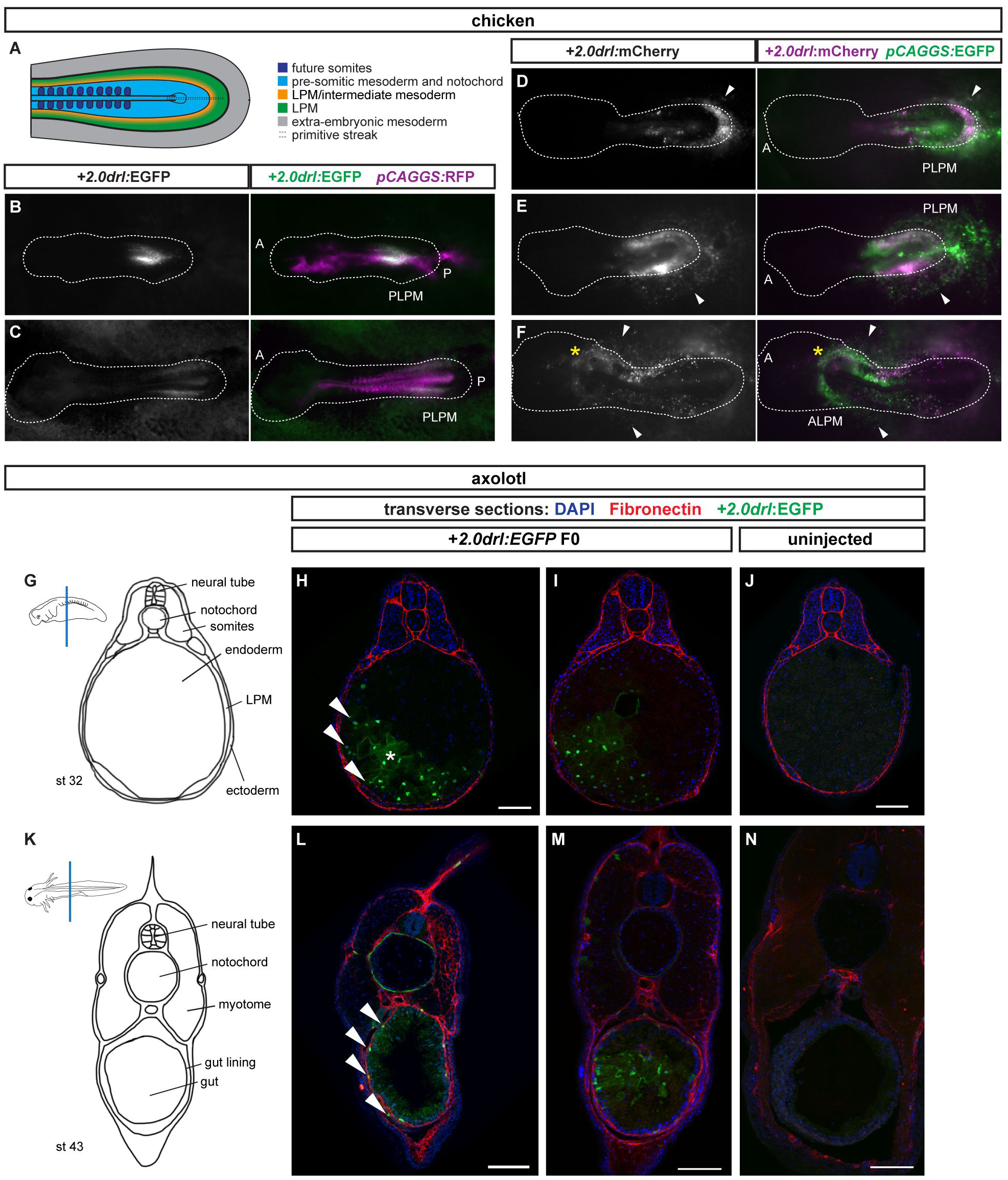
Examples of electroporated chicken embryos with +*2.0drl* reporters driving specific expression in the chicken LPM. (**A**) Posterior region of chicken embryo represented in a schematic showing different territories. (**B-F**) *Ex ovo* cultured chicken embryos electroporated with +*2.0drl cis*-regulatory reporters as (**B-C**) +*2.0drl:EGFP*, (**D-F**) +*2.0drl:mCherry* (magenta) together with control plasmid (**B-C**) *pCAGGS:RFP* (magenta) or (**D-G**) *pCAGGS:EGFP* at HH3+/HH4. Arrowheads indicate extra-embryonic endothelial/blood progenitors and asterisks the heart field (**F**). The dashed lines indicate the outline of the chicken embryos based on bright field imaging, with anterior (a) located to the left. (**G**) Schematic and (**H,I**) confocal Z-stack projections of a transverse section through the trunk region of the transgenic +2.0*drl*:EGFP and (**J**) negative control (uninjected) embryo counterstained with fibronectin antibody at tailbud stage 32, showing mesendoderm (arrowheads) and endodermal EGFP positive cells. (**K**) Schematic and (**L,M**) confocal Z-stack projections of transverse sections through the trunk region of the transgenic +2.0*drl*:EGFP and (**N**) negative control (uninjected) animals at stage 43 showing EGFP positive cells in the gut lining (arrowheads) consistent with LPM origin, and a few cells in the endoderm. Scale bars (**H-I, L-N**) 200µm.

**Fig. S10:**
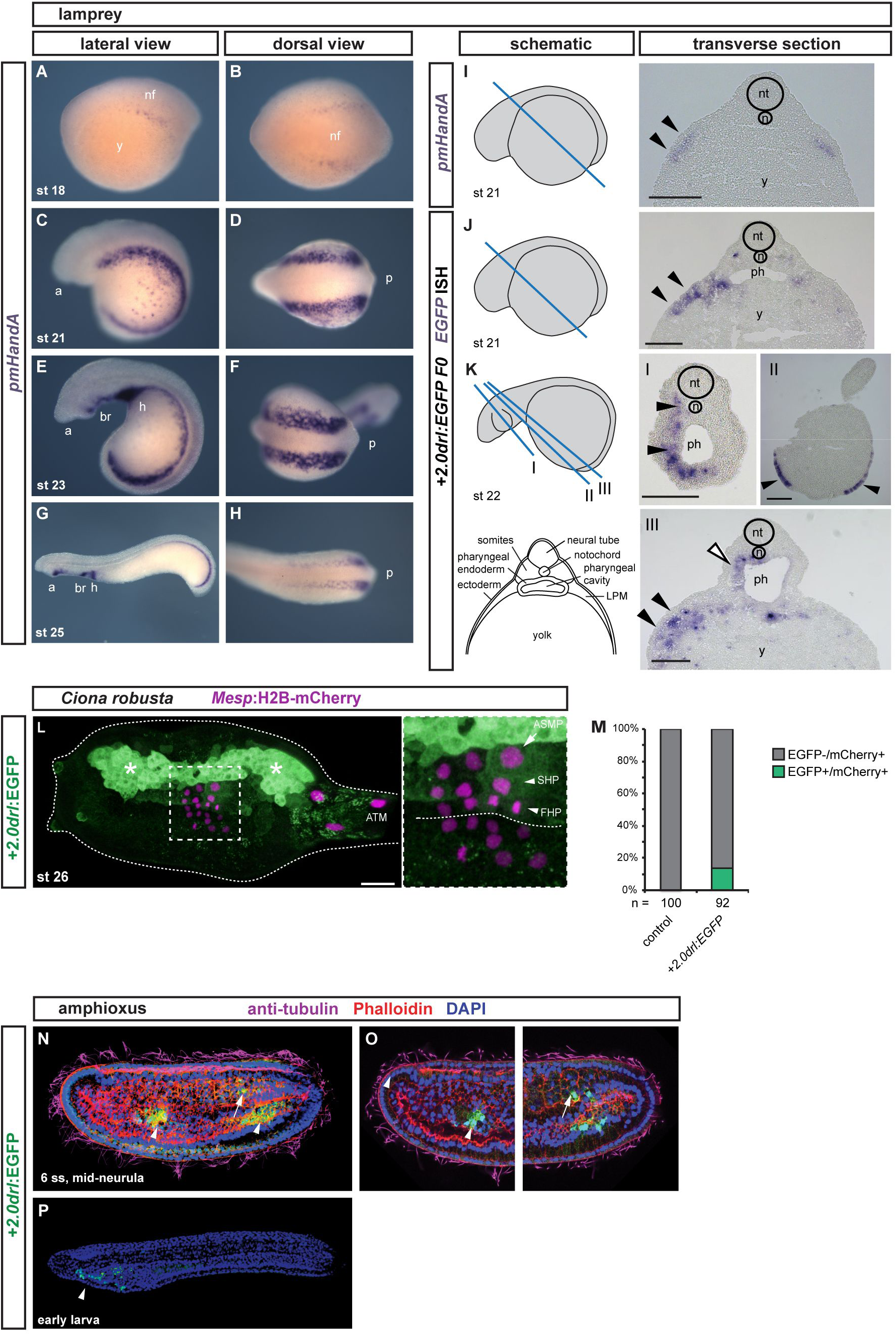
+*2.0drl* drives specific reporter expression in lamprey, *Ciona*, and amphioxus. (**A-D**) Whole-mount ISH for *pmHandA* marks the LPM in lamprey embryos from neurula to hatching stages (st 18-25). (**A**) Expression is first detectable as bilateral stripes in the PLPM at st 18. (**B**,**C**) Later, the PLPM expression expands covering the yolk, and *pmHandA* is upregulated in the heart tube, branchial region, and caudal to the heart. Lateral and dorsal views are shown with the anterior (a) of the embryo to the left. (**I-K**) Transverse sections of fixed st 21-22 embryos. The schematics indicate the plane of sectioning in blue and sections are shown with dorsal to the top. The schematic transverse section represents **K-III**. (**I**) ISH for *pmHandA* at st 21 marking the LPM (black arrowheads). (**J,K**) ISH for *EGFP* in transverse sections of +*2.0drl:EGFP* transient transgenic lamprey embryos fixed at st 21-22, showing enhancer activity in the anterior mesendoderm subjacent to the ectoderm (black arrowheads), in the pharyngeal endoderm (**K-III**, white arrowhead), in the pharyngeal mesoderm (**K**, black arrowheads), and in some embryos in the ectoderm as commonly observed unspecific expression of reporter plasmids (**K-II**, black arrowheads). (**L,M**) Immunostaining for EGFP in *Ciona* larvae embryo (st 26) expressing *drl* reporters (green) and *Mesp*:H2B-mCherry to track the B7.5 nuclei cell lineage (red). (**L**) Expression of +*2.0drl*-driven EGFP reporter in the larvae is stained in the mesenchymal lineage (white asterisks) and in the B7.5 cell progeny including ASM precursors (ASMP) (white arrow) and both cardiac first and second heart precursors (FHPs and SHPs) (white arrowheads), zoomed in box. Anterior to the left. (**M**) Proportion of larvae embryos expressing both GFP and mCherry in the B7.5 lineage when co-electroporated *drl* reporter and *Mesp*:H2B-mCherry in comparison to the control. n: number of electroporated larval halves. (**N-P**) Confocal Z-stack of amphioxus embryo at mid-neurula stage (6 ss), injected with +*2.0drl:EGFP*, showing specific reporter activity in the lateral endoderm (arrowhead), and ventral half of the somites (arrows). Embryos counterstained with Phalloidin (red), embryos/larvae with DAPI (blue), and anti-acetylated tubulin antibody (magenta). Lateral view shown as 3D-rendering (**N**) or Z-stack sagittal sections (**O**), anterior to the left and dorsal to the top. At early larvae stage (**P**), the activity of +*2.0drl* reporter was observed in the developing pharynx (n = 15/30, arrowhead). Branchial region (br), heart (h), neural folds (nf), neural tube (nt), notochord (n), pharynx (ph), and yolk (y) labeled (**I-K**). Scale bars (**I-K**) 200 µm, (**L**) 25 μm.

**Supplementary Movie legends:**

**Movie S1: 3D lightsheet imaging of the arising *drl* reporter-labeled LPM.**

SPIM time lapse imaging of a zebrafish embryo transgenic for *drl*:EGFP (green) together with the nuclear marker *actb2:*H2afva-mCherry (magenta) to visualize LPM emergence from 50% epiboly to 10 ss.

**Movie S2: Panoramic lightsheet imaging of the arising *drl* reporter-labeled LPM.**

Mercator projection of SPIM time lapse imaging of zebrafish embryos transgenic for *drl:EGFP* to visualize LPM emergence from 50% epiboly to 10 ss. From somitogenesis onwards, the embryo is oriented with anterior to the left and posterior to the right.

**Movie S3: 3D lightsheet imaging showing overlap between *drl* reporter expression and *sox17*-marked endoderm during gastrulation.**

SPIM time lapse imaging of *drl:mCherry* (red) combined with *sox17:EGFP* (green) to visualize the *drl*-expressing emerging cell population versus endoderm from 50% epiboly until 16 ss.

**Movie S4: Panoramic lightsheet imaging showing overlap between *drl* reporter expression and *sox17*-marked endoderm during gastrulation.**

Mercator projection of SPIM time lapse imaging of *drl:mCherry* (magenta) combined with *sox17:EGFP* to visualize the *drl*-expressing emerging cell population versus endoderm from 50% epiboly until 16 ss. From somitogenesis onwards, the embryo is oriented with anterior to the left and posterior to the right. Double-positive cells during gastrulation and early somitogenesis are shown in blue.

## References

Alev, C., Wu, Y., Nakaya, Y. and Sheng, G. (2013). Decoupling of amniote gastrulation and streak formation reveals a morphogenetic unity in vertebrate mesoderm induction. Development 140, 2691–6.

Alexander, J. and Stainier, D. Y. (1999). A molecular pathway leading to endoderm formation in zebrafish. Curr. Biol. 9, 1147–57.

Alexander, J., Rothenberg, M., Henry, G. L. and Stainier, D. Y. (1999). Casanova plays an early and essential role in endoderm formation in zebrafish. Dev. Biol. 215, 343–357.

Arnold, S. J., Hofmann, U. K., Bikoff, E. K. and Robertson, E. J. (2008). Pivotal roles for eomesodermin during axis formation, epithelium-to-mesenchyme transition and endoderm specification in the mouse. Development 135, 501–11.

Bailey, F. and Miller, A. (1921). Text-Book of Embryology. 4th ed. New York: William Wood and Co.

Bassett, A. R., Tibbit, C., Ponting, C. P. and Liu, J.-L. (2013). Highly efficient targeted mutagenesis of Drosophila with the CRISPR/Cas9 system. Cell Rep. 4, 220–8.

Becker, D., Eid, R. and Schughart, K. (1996). The limb/LPM enhancer of the murine Hoxb6 gene: reporter gene analysis in transgenic embryos and studies of DNA-protein interactions. Pharm. Acta Helv. 71, 29–35.

Beh, J., Shi, W., Levine, M., Davidson, B. and Christiaen, L. (2007). FoxF is essential for FGF-induced migration of heart progenitor cells in the ascidian Ciona intestinalis. Development 134, 3297–3305.

Bertrand, S., Escriva, H., Williams, N. A., Holland, N. D. and Holland, L. Z. (2011). Evolutionary crossroads in developmental biology: amphioxus. Development 138, 4819–30.

Bjornson, C. R. R., Griffin, K. J. P., Farr, G. H., Terashima, A., Himeda, C., Kikuchi, Y. and Kimelman, D. (2005). Eomesodermin Is a Localized Maternal Determinant Required for Endoderm Induction in Zebrafish. Dev. Cell 9, 523–533.

Bordzilovsakya, N. P., Dettlaf, T. A., Duhon, S. T. and Malacinski, G. M. (1989). Developmental-stage series of axolotl embryos. In Developmental Biology of the Axolotl (ed. Armstrong, J. B.) and Malacinski, G. M.), pp. 201–219. New York: Oxford University Press.

Bruce, A. E. E., Howley, C., Zhou, Y., Vickers, S. L., Silver, L. M., King, M. Lou and Ho, R. K. (2003). The maternally expressed zebrafish T-box gene eomesodermin regulates organizer formation. Development 130, 5503–5517.

Burger, A., Lindsay, H., Felker, A., Hess, C., Anders, C., Chiavacci, E., Zaugg, J., Weber, L. M. L. M., Catena, R., Jinek, M., et al. (2016). Maximizing mutagenesis with solubilized CRISPR-Cas9 ribonucleoprotein complexes. Development 143, 2025–37.

Chal, J. and Pourquié, O. (2017). Making muscle: skeletal myogenesis *in vivo* and *in vitro*. Development 144, 2104–2122.

Chandler, K. J., Chandler, R. L. and Mortlock, D. P. (2009). Identification of an ancient Bmp4 mesoderm enhancer located 46 kb from the promoter. Dev. Biol. 327, 590–602.

Charney, R. M., Forouzmand, E., Cho, J. S., Cheung, J., Paraiso, K. D., Yasuoka, Y., Takahashi, S., Taira, M., Blitz, I. L., Xie, X., et al. (2017). Foxh1 Occupies cis-Regulatory Modules Prior to Dynamic Transcription Factor Interactions Controlling the Mesendoderm Gene Program. Dev. Cell 40, 595–607.e4.

Chen, X., Rubock, M. J. and Whitman, M. (1996). A transcriptional partner for MAD proteins in TGF-β signalling. Nature 383, 691–696.

Chia, C. Y., Madrigal, P., Denil, S. L. I. J., Martinez, I., Garcia-Bernardo, J., El-Khairi, R., Chhatriwala, M., Shepherd, M. H., Hattersley, A. T., Dunn, N. R., et al. (2019). GATA6 Cooperates with EOMES/SMAD2/3 to Deploy the Gene Regulatory Network Governing Human Definitive Endoderm and Pancreas Formation. Stem Cell Reports 12, 57–70.

Chou, C.-W., Hsu, H.-C., You, M., Lin, J., Liu, Y.-W., Matsumoto, K., Yoshitomi, H., Rossant, J., Zaret, K. S., Nikolova, G., et al. (2016). The endoderm indirectly influences morphogenetic movements of the zebrafish head kidney through the posterior cardinal vein and VegfC. Sci. Rep. 6, 30677.

Costello, I., Pimeisl, I.-M., Dräger, S., Bikoff, E. K., Robertson, E. J. and Arnold, S. J. (2011). The T-box transcription factor Eomesodermin acts upstream of Mesp1 to specify cardiac mesoderm during mouse gastrulation. Nat. Cell Biol. 13, 1084–91.

DaCosta Byfield, S., Major, C., Laping, N. J. and Roberts, A. B. (2004). SB-505124 Is a Selective Inhibitor of Transforming Growth Factor-Type I Receptors ALK4, ALK5, and ALK7. Mol. Pharmacol. 65, 744–752.

Davidson, A. J. and Zon, L. I. (2004). The “definitive” (and ‘primitive’) guide to zebrafish hematopoiesis. Oncogene 23, 7233–7246.

Davidson, B., Shi, W. and Levine, M. (2005). Uncoupling heart cell specification and migration in the simple chordate Ciona intestinalis. Development 132, 4811–8.

De Los Angeles, A. and Daley, G. Q. (2013). Stem cells: Reprogramming in situ. Nature 502, 309–310.

Dick, A., Mayr, T., Bauer, H., Meier, A. and Hammerschmidt, M. (2000). Cloning and characterization of zebrafish smad2, smad3 and smad4. Gene 246, 69–80.

Dickmeis, T., Mourrain, P., Saint-Etienne, L., Fischer, N., Aanstad, P., Clark, M., Strähle, U. and Rosa, F. (2001). A crucial component of the endoderm formation pathway, CASANOVA, is encoded by a novel sox-related gene. Genes Dev. 15, 1487–1492.

Diogo, R., Kelly, R. G., Christiaen, L., Levine, M., Ziermann, J. M., Molnar, J. L., Noden, D. M. and Tzahor, E. (2015). A new heart for a new head in vertebrate cardiopharyngeal evolution. Nature 520, 466–73.

Du, S., Draper, B. W., Mione, M., Moens, C. B. and Bruce, A. (2012). Differential regulation of epiboly initiation and progression by zebrafish Eomesodermin A. Dev. Biol. 362, 11–23.

Dubrulle, J., Jordan, B. M., Akhmetova, L., Farrell, J. A., Kim, S.-H., Solnica-Krezel, L. and Schier, A. F. (2015). Response to Nodal morphogen gradient is determined by the kinetics of target gene induction. Elife 4, e05042.

Felker, A. and Mosimann, C. (2016). Contemporary zebrafish transgenesis with Tol2 and application for Cre/lox recombination experiments. Methods Cell Biol. 135, 219–44.

Felker, A., Nieuwenhuize, S., Dolbois, A., Blazkova, K., Hess, C., Low, L. W. L. W. L., Burger, S., Samson, N., Carney, T. J. T. J., Bartunek, P., et al. (2016). In Vivo Performance and Properties of Tamoxifen Metabolites for CreERT2 Control. PLoS One 11, e0152989.

Felker, A., Prummel, K. D. K. D., Merks, A. M. A. M., Mickoleit, M., Brombacher, E. C. E. C., Huisken, J., Panáková, D. and Mosimann, C. (2018). Continuous addition of progenitors forms the cardiac ventricle in zebrafish. Nat. Commun. 9, 2001.

Firulli, A. B., McFadden, D. G., Lin, Q., Srivastava, D. and Olson, E. N. (1998). Heart and extra-embryonic mesodermal defects in mouse embryos lacking the bHLH transcription factor Hand1. Nat. Genet. 18, 266–270.

Fuentes, M., Benito, E., Bertrand, S., Paris, M., Mignardot, A., Godoy, L., Jimenez-Delgado, S., Oliveri, D., Candiani, S., Hirsinger, E., et al. (2007). Insights into spawning behavior and development of the european amphioxus (Branchiostoma lanceolatum). J. Exp. Zool. Part B Mol. Dev. Evol. 308B, 484–493.

Gays, D., Hess, C., Camporeale, A., Ala, U., Provero, P., Mosimann, C. and Santoro, M. M. (2017). An exclusive cellular and molecular network governs intestinal smooth muscle cell differentiation in vertebrates. Development 144, 464–478.

Germain, S., Howell, M., Esslemont, G. M. and Hill, C. S. (2000). Homeodomain and winged-helix transcription factors recruit activated Smads to distinct promoter elements via a common Smad interaction motif. Genes Dev. 14, 435–51.

Gilbert, S. F. (2000). Developmental Biology, 6th edition.p. NBK10036. Sunderland, MA: Sinauer Associates, Inc.

Gritsman, K., Zhang, J., Cheng, S., Heckscher, E., Talbot, W. S. and Schier, A. F. (1999). The EGF-CFC Protein One-Eyed Pinhead Is Essential for Nodal Signaling. Cell 97, 121–132.

Gurdon, J. B. (1995). The Formation of Mesoderm and Muscle in Xenopus. In Organization of the Early Vertebrate Embryo, pp. 51–59. Boston, MA: Springer US.

Hagos, E. G. and Dougan, S. T. (2007). Time-dependent patterning of the mesoderm and endoderm by Nodal signals in zebrafish. BMC Dev. Biol. 7, 22.

Hatada, Y. and Stern, C. (1994). A fate map of the epiblast of the early chick embryo. Development 120, 2879–2889.

Helker, C. S. M., Schuermann, A., Pollmann, C., Chng, S. C., Kiefer, F., Reversade, B. and Herzog, W. (2015). The hormonal peptide Elabela guides angioblasts to the midline during vasculogenesis. Elife 4, e06726.

Henninger, J., Santoso, B., Hans, S., Durand, E., Moore, J., Mosimann, C., Brand, M., Traver, D. and Zon, L. (2017). Clonal fate mapping quantifies the number of haematopoietic stem cells that arise during development. Nat. Cell Biol. 19,.

Henry, G. L. and Melton, D. A. (1998). Mixer, a homeobox gene required for endoderm development. Science 281, 91–6.

Herbomel, P., Thisse, B. and Thisse, C. (1999). Ontogeny and behaviour of early macrophages in the zebrafish embryo. Development 126, 3735–3745.

Hild, M., Dick, A., Rauch, G. J., Meier, A., Bouwmeester, T., Haffter, P. and Hammerschmidt, M. (1999). The smad5 mutation somitabun blocks Bmp2b signaling during early dorsoventral patterning of the zebrafish embryo. Development 126, 2149–59.

Hill, C. S. (2018). Spatial and temporal control of NODAL signaling. Curr. Opin. Cell Biol. 51, 50–57.

Hockman, D., Burns, A. A. J., Schlosser, G., Gates, K. P. K. P., Jevans, B., Mongera, A., Fisher, S., Unlu, G., Knapik, E. W. E. W., Kaufman, C. K. C. K., et al. (2017). Evolution of the hypoxia-sensitive cells involved in amniote respiratory reflexes. Elife 6, e21231.

Holland, N. D. (2018). Formation of the initial kidney and mouth opening in larval amphioxus studied with serial blockface scanning electron microscopy (SBSEM). Evodevo 9, 16.

Holland, N. D., Venkatesh, T. V, Holland, L. Z., Jacobs, D. K. and Bodmer, R. (2003). AmphiNk2-tin, an amphioxus homeobox gene expressed in myocardial progenitors: insights into evolution of the vertebrate heart. Dev. Biol. 255, 128–37.

Hotta, K., Mitsuhara, K., Takahashi, H., Inaba, K., Oka, K., Gojobori, T. and Ikeo, K. (2007). A web-based interactive developmental table for the ascidian *Ciona intestinalis*, including 3D real-image embryo reconstructions: I. From fertilized egg to hatching larva. Dev. Dyn. 236, 1790–1805.

Ieda, M., Fu, J.-D., Delgado-Olguin, P., Vedantham, V., Hayashi, Y., Bruneau, B. G., Srivastava, D., Kamp, T. J., Reinecke, H., Murry, C. E., et al. (2010). Direct reprogramming of fibroblasts into functional cardiomyocytes by defined factors. Cell 142, 375–86.

Jin, H., Xu, J., Qian, F., Du, L., Tan, C. Y., Lin, Z., Peng, J. and Wen, Z. (2006). The 5’ zebrafish scl promoter targets transcription to the brain, spinal cord, and hematopoietic and endothelial progenitors. Dev Dyn 235, 60–67.

Kaplan, N., Razy-Krajka, F. and Christiaen, L. (2015). Regulation and evolution of cardiopharyngeal cell identity and behavior: insights from simple chordates. Curr. Opin. Genet. Dev. 32, 119–128.

Khan, A., Fornes, O., Stigliani, A., Gheorghe, M., Castro-Mondragon, J. A., van der Lee, R., Bessy, A., Chèneby, J., Kulkarni, S. R., Tan, G., et al. (2018). JASPAR 2018: update of the open-access database of transcription factor binding profiles and its web framework. Nucleic Acids Res. 46, D260–D266.

Khattak, S., Murawala, P., Andreas, H., Kappert, V., Schuez, M., Sandoval-Guzmán, T., Crawford, K. and Tanaka, E. M. (2014). Optimized axolotl (Ambystoma mexicanum) husbandry, breeding, metamorphosis, transgenesis and tamoxifen-mediated recombination. Nat. Protoc. 9, 529–540.

Kikuchi, Y., Trinh, L. A., Reiter, J. F., Alexander, J., Yelon, D. and Stainier, D. Y. (2000). The zebrafish bonnie and clyde gene encodes a Mix family homeodomain protein that regulates the generation of endodermal precursors. Genes Dev. 14, 1279–89.

Kikuchi, Y., Agathon, A., Alexander, J., Thisse, C., Waldron, S., Yelon, D., Thisse, B. and Stainier, D. Y. (2001). casanova encodes a novel Sox-related protein necessary and sufficient for early endoderm formation in zebrafish. Genes Dev. 15, 1493–1505.

Kozmik, Z., Holland, L. Z., Schubert, M., Lacalli, T. C., Kreslova, J., Vlcek, C. and Holland, N. D. (2001). Characterization of amphioxus Amphivent, an evolutionarily conserved marker for chordate ventral mesoderm. Genesis 29, 172–179.

Kozmikova, I. and Kozmik, Z. (2015). Gene regulation in amphioxus: An insight from transgenic studies in amphioxus and vertebrates. Mar. Genomics 24, 159–166.

Krens, S. F. G., Möllmert, S. and Heisenberg, C.-P. (2011). Enveloping cell-layer differentiation at the surface of zebrafish germ-layer tissue explants. Proc. Natl. Acad. Sci. U. S. A. 108, E9–10; author reply E11.

Kunwar, P. S., Zimmerman, S., Bennett, J. T., Chen, Y., Whitman, M. and Schier, A. F. (2003). Mixer/Bon and FoxH1/Sur have overlapping and divergent roles in Nodal signaling and mesendoderm induction. Development 130, 5589–99.

Kuraku, S., Takio, Y., Sugahara, F., Takechi, M. and Kuratani, S. (2010). Evolution of oropharyngeal patterning mechanisms involving Dlx and endothelins in vertebrates. Dev. Biol. 341, 315–23.

Kusakabe, R. and Kuratani, S. (2007). Evolutionary perspectives from development of mesodermal components in the lamprey. Dev. Dyn. 236, 2410–2420.

Kwan, K. M., Fujimoto, E., Grabher, C., Mangum, B. D., Hardy, M. E., Campbell, D. S., Parant, J. M., Yost, H. J., Kanki, J. P. and Chien, C. Bin (2007). The Tol2kit: a multisite gateway-based construction kit for Tol2 transposon transgenesis constructs. Dev Dyn 236, 3088–3099.

Langdon, Y. G. and Mullins, M. C. (2011). Maternal and zygotic control of zebrafish dorsoventral axial patterning. Annu Rev Genet 45, 357–377.

Lee, J. H., Protze, S. I., Laksman, Z., Backx, P. H. and Keller, G. M. (2017). Human Pluripotent Stem Cell-Derived Atrial and Ventricular Cardiomyocytes Develop from Distinct Mesoderm Populations. Cell Stem Cell 21, 179–194.e4.

Lindsay, H., Burger, A., Biyong, B., Felker, A., Hess, C., Zaugg, J., Chiavacci, E., Anders, C., Jinek, M., Mosimann, C., et al. (2016). CrispRVariants charts the mutation spectrum of genome engineering experiments. Nat. Biotechnol. 34, 701–702.

Martin, J. F. and Olson, E. N. (2000). Identification of a prx1 limb enhancer. genesis 26, 225–229.

Massagué, J. (2012). TGFβ signalling in context. Nat. Rev. Mol. Cell Biol. 13, 616–630.

McDole, K., Guignard, L., Amat, F., Berger, A., Malandain, G., Royer, L. A., Turaga, S. C., Branson, K. and Keller, P. J. (2018). In Toto Imaging and Reconstruction of Post-Implantation Mouse Development at the Single-Cell Level. Cell 175, 859–876.e33.

McEwen, G. K., Goode, D. K., Parker, H. J., Woolfe, A., Callaway, H. and Elgar, G. (2009). Early evolution of conserved regulatory sequences associated with development in vertebrates. PLoS Genet. 5, e1000762.

Mead, P. E., Brivanlou, I. H., Kelley, C. M. and Zon, L. I. (1996). BMP-4-responsive regulation of dorsal–ventral patterning by the homeobox protein Mix.1. Nature 382, 357–360.

Mendjan, S., Mascetti, V. L., Ortmann, D., Ortiz, M., Karjosukarso, D. W., Ng, Y., Moreau, T. and Pedersen, R. A. (2014). NANOG and CDX2 Pattern Distinct Subtypes of Human Mesoderm during Exit from Pluripotency. Cell Stem Cell 15, 310–25.

Miyazono, K.-I., Moriwaki, S., Ito, T., Kurisaki, A., Asashima, M. and Tanokura, M. (2018). Hydrophobic patches on SMAD2 and SMAD3 determine selective binding to cofactors. Sci. Signal. 11, eaao7227.

Moreau, C., Caldarelli, P., Rocancourt, D., Roussel, J., Denans, N., Pourquie, O. and Gros, J. (2019). Timed Collinear Activation of Hox Genes during Gastrulation Controls the Avian Forelimb Position. Curr. Biol. 29, 35–50.e4.

Mosimann, C., Kaufman, C. K., Li, P., Pugach, E. K. E. K., Tamplin, O. J. O. J. and Zon, L. I. L. I. (2011). Ubiquitous transgene expression and Cre-based recombination driven by the ubiquitin promoter in zebrafish. Development 138, 169–177.

Mosimann, C., Panáková, D., Werdich, A. A. A., Musso, G., Burger, A., Lawson, K. L. K. L., Carr, L. A. L. A., Nevis, K. R. K. R., Sabeh, M. K. K., Zhou, Y., et al. (2015). Chamber identity programs drive early functional partitioning of the heart. Nat. Commun. 6, 8146.

Mullins, M. C., Hammerschmidt, M., Kane, D. A., Odenthal, J., Brand, M., van Eeden, F. J., Furutani-Seiki, M., Granato, M., Haffter, P., Heisenberg, C. P., et al. (1996). Genes establishing dorsoventral pattern formation in the zebrafish embryo: the ventral specifying genes. Development 123, 81–93.

Murry, C. E. and Keller, G. (2008). Differentiation of embryonic stem cells to clinically relevant populations: lessons from embryonic development. Cell 132, 661–80.

Nelson, A. C., Cutty, S. J., Niini, M., Stemple, D. L., Flicek, P., Houart, C., Bruce, A. and Wardle, F. C. (2014). Global identification of Smad2 and Eomesodermin targets in zebrafish identifies a conserved transcriptional network in mesendoderm and a novel role for Eomesodermin in repression of ectodermal gene expression. BMC Biol. 12, 81.

Nelson, A. C., Cutty, S. J., Gasiunas, S. N., Deplae, I., Stemple, D. L. and Wardle, F. C. (2017). In Vivo Regulation of the Zebrafish Endoderm Progenitor Niche by T-Box Transcription Factors. Cell Rep. 19, 2782–2795.

New, D. A. T. (1955). A New Technique for the Cultivation of the Chick Embryo in vitro. Development 3, 326–331.

Onimaru, K., Shoguchi, E., Kuratani, S. and Tanaka, M. (2011). Development and evolution of the lateral plate mesoderm: comparative analysis of amphioxus and lamprey with implications for the acquisition of paired fins. Dev. Biol. 359, 124–36.

Ormestad, M., Astorga, J. and Carlsson, P. (2004). Differences in the embryonic expression patterns of mouse Foxf1 and −2 match their distinct mutant phenotypes. Dev. Dyn. 229, 328–33.

Osterwalder, M., Speziale, D., Shoukry, M., Mohan, R., Ivanek, R., Kohler, M., Beisel, C., Wen, X., Scales, S. J., Christoffels, V. M., et al. (2014). HAND2 targets define a network of transcriptional regulators that compartmentalize the early limb bud mesenchyme. Dev. Cell 31, 345–57.

Parker, H. J., Piccinelli, P., Sauka-Spengler, T., Bronner, M. and Elgar, G. (2011). Ancient Pbx-Hox signatures define hundreds of vertebrate developmental enhancers. BMC Genomics 12, 637.

Parker, H. J., Sauka-Spengler, T., Bronner, M. and Elgar, G. (2014a). A reporter assay in lamprey embryos reveals both functional conservation and elaboration of vertebrate enhancers. PLoS One 9, e85492.

Parker, H. J., Bronner, M. E. and Krumlauf, R. (2014b). A Hox regulatory network of hindbrain segmentation is conserved to the base of vertebrates. Nature 514, 490–493.

Pascual-Anaya, J., Albuixech-Crespo, B., Somorjai, I. M. L., Carmona, R., Oisi, Y., Álvarez, S., Kuratani, S., Muñoz-Chápuli, R. and Garcia-Fernàndez, J. (2013). The evolutionary origins of chordate hematopoiesis and vertebrate endothelia. Dev. Biol. 375, 182–192.

Perens, E. A., Garavito-Aguilar, Z. V, Guio-Vega, G. P., Peña, K. T., Schindler, Y. L. and Yelon, D. (2016). Hand2 inhibits kidney specification while promoting vein formation within the posterior mesoderm. Elife 5, e19941.

Pfeiffer, M. J., Quaranta, R., Piccini, I., Fell, J., Rao, J., Röpke, A., Seebohm, G. and Greber, B. (2018). Cardiogenic programming of human pluripotent stem cells by dose-controlled activation of EOMES. Nat. Commun. 9, 440.

Picker, A., Scholpp, S., Böhli, H., Takeda, H. and Brand, M. (2002). A novel positive transcriptional feedback loop in midbrain-hindbrain boundary development is revealed through analysis of the zebrafish pax2.1 promoter in transgenic lines. Development 129, 3227–39.

Poulain, M. and Lepage, T. (2002). Mezzo, a paired-like homeobox protein is an immediate target of Nodal signalling and regulates endoderm specification in zebrafish. Development 129,.

Racioppi, C., Kamal, A. K., Razy-Krajka, F., Gambardella, G., Zanetti, L., di Bernardo, D., Sanges, R., Christiaen, L. A. and Ristoratore, F. (2014). Fibroblast growth factor signalling controls nervous system patterning and pigment cell formation in Ciona intestinalis. Nat. Commun. 5, 4830.

Racioppi, C., Wiechecki, K. A. and Christiaen, L. (2019). Combinatorial chromatin dynamics foster accurate cardiopharyngeal fate choices. bioRxiv 546945.

Reichenbach, B., Delalande, J.-M., Kolmogorova, E., Prier, A., Nguyen, T., Smith, C. M., Holzschuh, J. and Shepherd, I. T. (2008). Endoderm-derived Sonic hedgehog and mesoderm Hand2 expression are required for enteric nervous system development in zebrafish. Dev. Biol. 318, 52–64.

Rogers, K. W. and Müller, P. (2019). Nodal and BMP dispersal during early zebrafish development. Dev. Biol. 447, 14–23.

Rogers, K. W., Lord, N. D., Gagnon, J. A., Pauli, A., Zimmerman, S., Aksel, D. C., Reyon, D., Tsai, S. Q., Joung, J. K. and Schier, A. F. (2017). Nodal patterning without Lefty inhibitory feedback is functional but fragile. Elife 6, e28785.

Rojas, A., De Val, S., Heidt, A. B., Xu, S.-M., Bristow, J. and Black, B. L. (2005). Gata4 expression in lateral mesoderm is downstream of BMP4 and is activated directly by Forkhead and GATA transcription factors through a distal enhancer element. Development 132, 3405–17.

Sakaguchi, T., Kuroiwa, A. and Takeda, H. (2001). A novel sox gene, 226D7, acts downstream of Nodal signaling to specify endoderm precursors in zebrafish. Mech. Dev. 107, 25–38.

Sakaguchi, T., Kikuchi, Y., Kuroiwa, A., Takeda, H. and Stainier, D. Y. R. (2006). The yolk syncytial layer regulates myocardial migration by influencing extracellular matrix assembly in zebrafish. Development 133, 4063–4072.

Sánchez-Iranzo, H., Galardi-Castilla, M., Minguillón, C., Sanz-Morejón, A., González-Rosa, J. M. J. M., Felker, A., Ernst, A., Guzmán-Martínez, G., Mosimann, C. and Mercader, N. (2018). Tbx5a lineage tracing shows cardiomyocyte plasticity during zebrafish heart regeneration. Nat. Commun. 9, 428.

Sauka-Spengler, T., Meulemans, D., Jones, M. and Bronner-Fraser, M. (2007). Ancient Evolutionary Origin of the Neural Crest Gene Regulatory Network. Dev. Cell 13, 405–420.

Schier, a F., Neuhauss, S. C., Helde, K. a, Talbot, W. S. and Driever, W. (1997). The one-eyed pinhead gene functions in mesoderm and endoderm formation in zebrafish and interacts with no tail. Development 124, 327–42.

Schindelin, J., Arganda-Carreras, I., Frise, E., Kaynig, V., Longair, M., Pietzsch, T., Preibisch, S., Rueden, C., Saalfeld, S., Schmid, B., et al. (2012). Fiji: an open-source platform for biological-image analysis. Nat. Methods 9, 676–82.

Schmid, B., Shah, G., Scherf, N., Weber, M., Thierbach, K., Campos, C. P., Roeder, I., Aanstad, P. and Huisken, J. (2013). High-speed panoramic light-sheet microscopy reveals global endodermal cell dynamics. Nat Commun 4, 2207.

Shimeld, S. M. and Donoghue, P. C. J. (2012). Evolutionary crossroads in developmental biology: cyclostomes (lamprey and hagfish). Development 139, 2091–9.

Slagle, C. E., Aoki, T. and Burdine, R. D. (2011). Nodal-dependent mesendoderm specification requires the combinatorial activities of FoxH1 and Eomesodermin. PLoS Genet. 7, e1002072.

Slukvin II (2013). Hematopoietic specification from human pluripotent stem cells: current advances and challenges toward de novo generation of hematopoietic stem cells. Blood 122, 4035–4046.

Song, K., Nam, Y.-J., Luo, X., Qi, X., Tan, W., Huang, G. N., Acharya, A., Smith, C. L., Tallquist, M. D., Neilson, E. G., et al. (2012). Heart repair by reprogramming non-myocytes with cardiac transcription factors. Nature 485, 599–604.

Stainier, D. Y., Fouquet, B., Chen, J. N., Warren, K. S., Weinstein, B. M., Meiler, S. E., Mohideen, M. A., Neuhauss, S. C., Solnica-Krezel, L., Schier, A. F., et al. (1996). Mutations affecting the formation and function of the cardiovascular system in the zebrafish embryo. Development 123, 285–92.

Stolfi, A. and Christiaen, L. (2012). Genetic and genomic toolbox of the chordate Ciona intestinalis. Genetics 192, 55–66.

Takasato, M. and Little, M. H. (2015). The origin of the mammalian kidney: implications for recreating the kidney in vitro. Development 142, 1937–1947.

Takasato, M., Er, P. X., Becroft, M., Vanslambrouck, J. M., Stanley, E. G., Elefanty, A. G. and Little, M. H. (2014). Directing human embryonic stem cell differentiation towards a renal lineage generates a self-organizing kidney. Nat. Cell Biol. 16, 118–26.

Takeuchi, J. K. and Bruneau, B. G. (2009). Directed transdifferentiation of mouse mesoderm to heart tissue by defined factors. Nature 459, 708–11.

Tamplin, O. J., Durand, E. M., Carr, L. A., Childs, S. J., Hagedorn, E. J., Li, P., Yzaguirre, A. D., Speck, N. A. and Zon, L. I. (2015). Hematopoietic stem cell arrival triggers dynamic remodeling of the perivascular niche. Cell 160, 241–52.

Technau, U. and Scholz, C. B. (2003). Origin and evolution of endoderm and mesoderm. Int. J. Dev. Biol. 47, 531–539.

Thisse, C. and Thisse, B. (2008). High-resolution in situ hybridization to whole-mount zebrafish embryos. Nat. Protoc. 3, 59–69.

Tonegawa, A., Funayama, N., Ueno, N. and Takahashi, Y. (1997). Mesodermal subdivision along the mediolateral axis in chicken controlled by different concentrations of BMP-4. Development 124, 1975–1984.

Warga, R. M. and Nüsslein-Volhard, C. (1999). Origin and development of the zebrafish endoderm. Development 126, 827–38.

Westerfield, M. (2007). The Zebrafish Book: a guide for the laboratory use of zebrafish (Danio rerio). 5th ed. Eugene: University of Oregon Press.

Whitman, M. (2001). Nodal Signaling in Early Vertebrate Embryos: Themes and Variations. Dev. Cell 1, 605–617.

Xu, P., Zhu, G., Wang, Y., Sun, J., Liu, X., Chen, Y.-G. and Meng, A. (2014). Maternal Eomesodermin regulates zygotic nodal gene expression for mesendoderm induction in zebrafish embryos. J. Mol. Cell Biol. 6, 272–285.

Yabe, T., Hoshijima, K., Yamamoto, T. and Takada, S. (2016). Quadruple zebrafish mutant reveals different roles of Mesp genes in somite segmentation between mouse and zebrafish. Development 143, 2842–2852.

Yelon, D., Ticho, B., Halpern, M. E., Ruvinsky, I., Ho, R. K., Silver, L. M. and Stainier, D. Y. R. (2000). The bHLH transcription factor hand2 plays parallel roles in zebrafish heart and pectoral fin development. Development 127, 2573–82.

Yin, C., Kikuchi, K., Hochgreb, T., Poss, K. D. and Stainier, D. Y. R. (2010). Hand2 regulates extracellular matrix remodeling essential for gut-looping morphogenesis in zebrafish. Dev. Cell 18, 973–984.

Yu, P. B., Hong, C. C., Sachidanandan, C., Babitt, J. L., Deng, D. Y., Hoyng, S. A., Lin, H. Y., Bloch, K. D. and Peterson, R. T. (2008). Dorsomorphin inhibits BMP signals required for embryogenesis and iron metabolism. Nat. Chem. Biol. 4, 33–41.

Zhang, X. Y. and Rodaway, A. R. F. (2007). SCL-GFP transgenic zebrafish: In vivo imaging of blood and endothelial development and identification of the initial site of definitive hematopoiesis. Dev. Biol. 307, 179–194.

Zhang, H., Fraser, S. T., Papazoglu, C., Hoatlin, M. E. and Baron, M. H. (2009). Transcriptional activation by the Mixl1 homeodomain protein in differentiating mouse embryonic stem cells. Stem Cells 27, 2884–95.

Zhu, H., Traver, D., Davidson, A. J., Dibiase, A., Thisse, C., Thisse, B., Nimer, S. and Zon, L. I. (2005). Regulation of the lmo2 promoter during hematopoietic and vascular development in zebrafish. Dev Biol 281, 256–269.

